# Active Dissociation of Intracortical Spiking and High Gamma Activity

**DOI:** 10.1101/2025.07.10.663559

**Authors:** Tianhao Lei, Michael R Scheid, Robert D Flint, Joshua I Glaser, Marc W Slutzky

## Abstract

Cortical high gamma activity (HGA) is used in many scientific investigations, yet its biophysical source is a matter of debate. Two leading hypotheses are that HGA predominantly represents summed postsynaptic potentials or–more commonly– predominantly represents summed local spikes. If the latter were true, the nearest neurons to an electrode should contribute most to HGA recorded on that electrode. We trained subjects to decouple local spiking from HGA on a single electrode using a brain-machine interface. Their ability to decouple them suggested that HGA is likely not simply generated by summed local spiking. Instead, HGA correlated with neuronal population co-firing of neurons that were widely distributed across millimeters. The neuronal spikes that contributed more to this co-firing also contributed more to, and preceded, spike-triggered HGA. These results suggest that HGA arises predominantly from summed postsynaptic potentials triggered by synchronous co-firing of widely distributed neurons.

## MAIN

High gamma band activity (HGA, power in the 70-300 Hz band of the local field potential (LFP), sometimes called broadband activity^1,2^) is a highly informative cortical signal that has been used to investigate a range of topics including sensory processing^3,4^, motor control^5–8^, brain-machine interfaces (BMIs)^9–12^, and cognitive neuroscience, including attention, memory, language, and mood^13–19^. These diverse applications underscore the significance of HGA as a robust and informative measure of mesoscale neural processing with high spatiotemporal resolution^20^.

Despite this wide range of usage, the origin of HGA remains rather controversial. A number of studies have suggested HGA is an “index” of *local* neuronal action potentials (spikes), as either capturing the total firing rate^1,2,20–28^, or synchronous firing of neurons in local populations^18,29–32^. Two seminal studies found that HGA was modestly correlated with local spiking activity near the recording electrode^29,31^. Other studies suggested that correlation between HGA and spikes was due to spectral power from spikes “leaking” into the high gamma band range^33–37^. Many subsequent studies have cited these findings in equating HGA with (an index of) summed local population spiking activity, despite the low correlations with individual neurons.

However, other results suggested that HGA is not simply an index of local spiking activity. Two studies found low correlation between spikes and HGA recorded from the same electrode^16,38^. Rather, HGA correlated more with low-dimensional neuronal population spike rates from a more distributed cortical area (16 mm^2^) than with spiking from single electrodes^38^. The mechanism of this higher correlation with population spiking activity is unknown; volume conduction is not the explanation at such a long range. Evidence that HGA is largely generated by postsynaptic potentials^33^ may indirectly support this finding. For example, HGA in layer 5 of visual cortex was correlated with spiking, while HGA in layers 2-3 was not correlated with spiking^39^. Further, blocking NMDA activity silenced HGA but not local spiking, which the authors found consistent with HGA generation by postsynaptic potentials.

Thus, there is still substantial disagreement regarding the biophysical origin of HGA and its relationship with spiking. To summarize the literature, there are two hypotheses about the main biophysical source of HGA (Fig. 1): (1) HGA originates predominantly from summed waveforms of nearby action potentials via volume conduction (“leakage”), and (2) HGA originates predominantly from summed postsynaptic activity. The second hypothesis implies that HGA will be highest when inputs to the local neuronal population fire synchronously. Note that these two hypotheses do not necessarily mean that the origin is exclusively one or the other, and there could be some contribution of both to HGA, as some studies imply; however, the broader literature tends to attribute the biophysical source to one or the other. Further, the vast majority of prior evidence about HGA’s origins was based on correlations or modeling.

**Figure 1.**
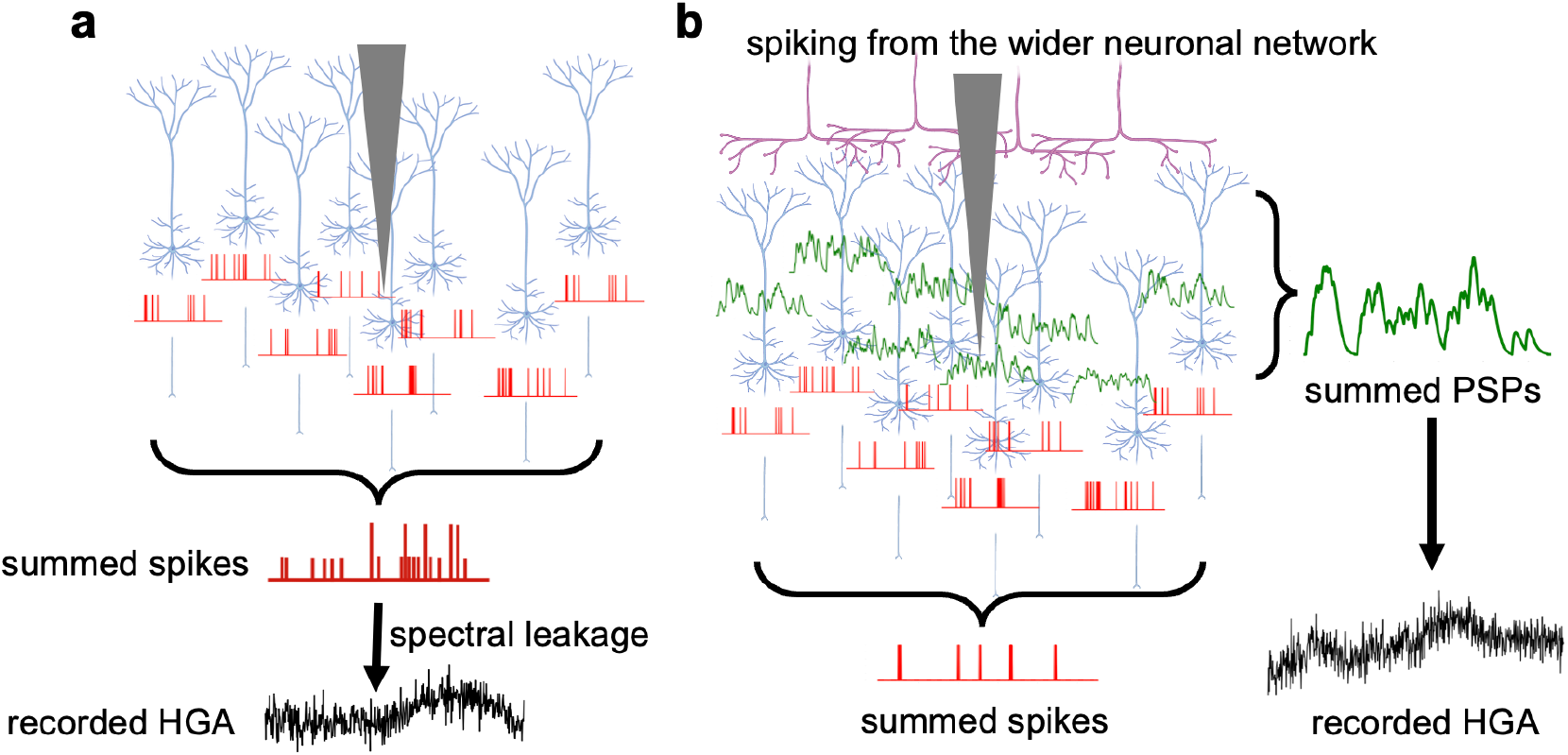
Illustration of potential sources of HGA generation. **a**, HGA recorded by an electrode (vertical triangle) predominantly originates from spectral leakage of summed nearby spikes through volume conduction. **b**, Recorded HGA predominantly reflects summed postsynaptic potentials (PSPs) that are triggered by spiking activity across the network that trigger PSPs near the recording electrode. Red traces are spike trains, at or near the axon hillock. Green traces are PSPs in dendrites and cell bodies. Purple neurites represent incoming axons from more distantly located neurons.

Here, we developed a novel brain machine interface (BMI) paradigm to causally test these hypotheses. This test required monkeys to modulate HGA independently from spike rate on a single electrode (Fig. 2). We found that monkeys could indeed decouple spiking from HGA on the same electrode in the primary motor cortex (M1) within a few sessions. Further, we investigated how HGA relates to the neural population activity across an area of 16 mm^2^. HGA on the control electrode did not exclusively correlate with action potentials from nearby neurons. Rather, HGA was correlated with low-dimensional, synchronous neuronal population firing (“co-firing”^8^) patterns spread across the array. We corroborated this result by using spike-triggered averages of HGA to determine that spikes throughout the network preceded local HGA by a similar lag to that between presynaptic spikes and postsynaptic potentials. Finally, we explored how the neuronal network activity could explain independent modulation of HGA and spike rate. Together, these results provide evidence consistent with the predominant source of HGA being primarily summed nearby postsynaptic activity triggered by synchronous firing activity throughout the wider neuronal ensemble. These findings help reconcile prior results showing highly correlated averaged HGA and spiking across the array with those supporting a postsynaptic potential origin of HGA.

**Figure 2.**
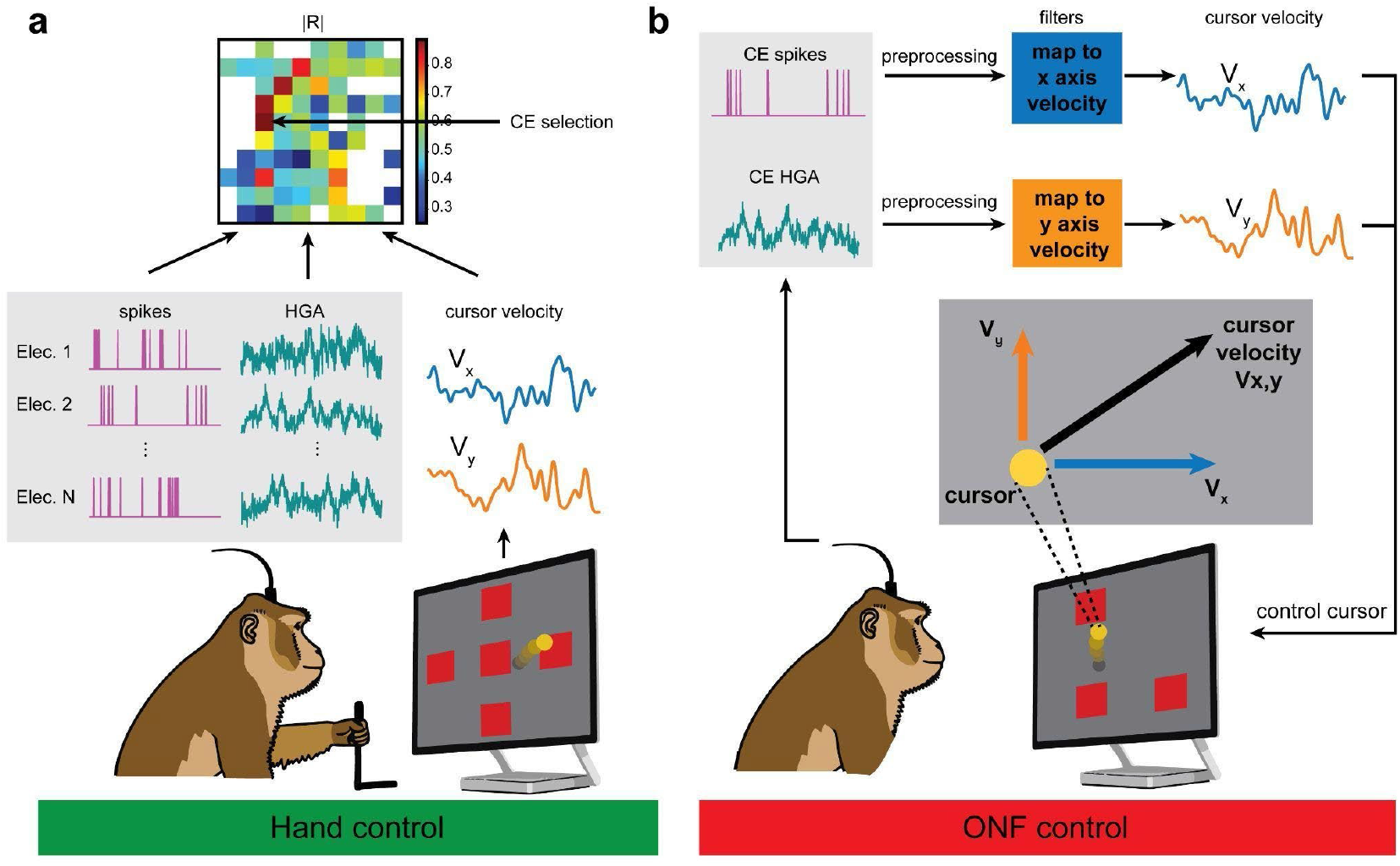
Task schematic. **a**, An illustration of the hand control task and the selection of control electrodes (CEs). The monkey used a manipulandum to move a cursor in a 4-target center-out reaching task. Spike rates and cursor velocity were simultaneously recorded. The absolute value of correlation (|R|, see Methods) between the two signals on each electrode was calculated and used to select the CE. See Methods. **b**, An illustration of the 2D ONF control task. The monkey was required to control the cursor velocity with HGA and spike rate using pre-trained filters with fixed weights from the CE. The task was a 2-target center-out task.

## RESULTS

### Monkeys perform well on a BMI task requiring dissociation of spikes from HGA

We designed an orthogonal neurofeedback (ONF) BMI to test whether monkeys could dissociate spiking rates from HGA recorded on the same electrode. Three monkeys first performed a 2D center-out cursor control task using a manipulandum (“hand control”; see Methods). They then performed the ONF task, which mapped unsorted spikes and HGA from an individual control electrode (CE) to orthogonal components of cursor velocity (Fig. 2). This paradigm, which we have used with myoelectric control in humans^40–44^, required the monkeys to dissociate HGA from nearby spikes in order to reach targets on the cardinal axes. For each monkey, we selected three electrodes with relatively high spike-HGA correlation as CEs for the task (Methods). All monkeys performed at least one 10-minute file of hand control at the beginning as well as at the end of the same day. Numbers of sessions of recordings for each monkey-CE combination are summarized in Extended Data Table 1.

Monkeys were able to perform the task with high proficiency, often starting in the first session (Fig. 3). In general, there were a considerable number of failed trials very early in ONF training, and the trajectories were somewhat more variable. The number of failed trials quickly decreased and trajectories became more consistent (Fig. 3a, Extended Data Fig. 1a). We quantified behavioral performance across all CEs for all sessions (see Methods). The monkeys’ performance largely started high and improved slightly, as measured by increased success rate (two-sided Wilcoxon rank-sum test, p=0.01; Fig. 3b), decreased time to target (two-sided Wilcoxon rank-sum test, p=0.04; Fig. 3c), and decreased path length (two-sided Wilcoxon rank-sum test, p=0.0001; Fig. 3d). However, we did not observe a significant change in takeoff or entry angles (one-sided one sample t-test, takeoff angle: p=0.14; entry angle: 0.053, see Methods; Fig. 3e,f). To examine these performance measures more closely, we removed the first session from these metrics; in the remaining sessions, we did not observe a significant improvement in success rate (p=0.87) or time to target (p=0.17). In fact, the performance improvement was strongest within the first 50 trials, then plateaued (Extended Data Fig. 1b). Combined with the consistent takeoff angle and entry angle across sessions, these results suggested that monkeys can innately modulate the two signals independently.

**Figure 3.**
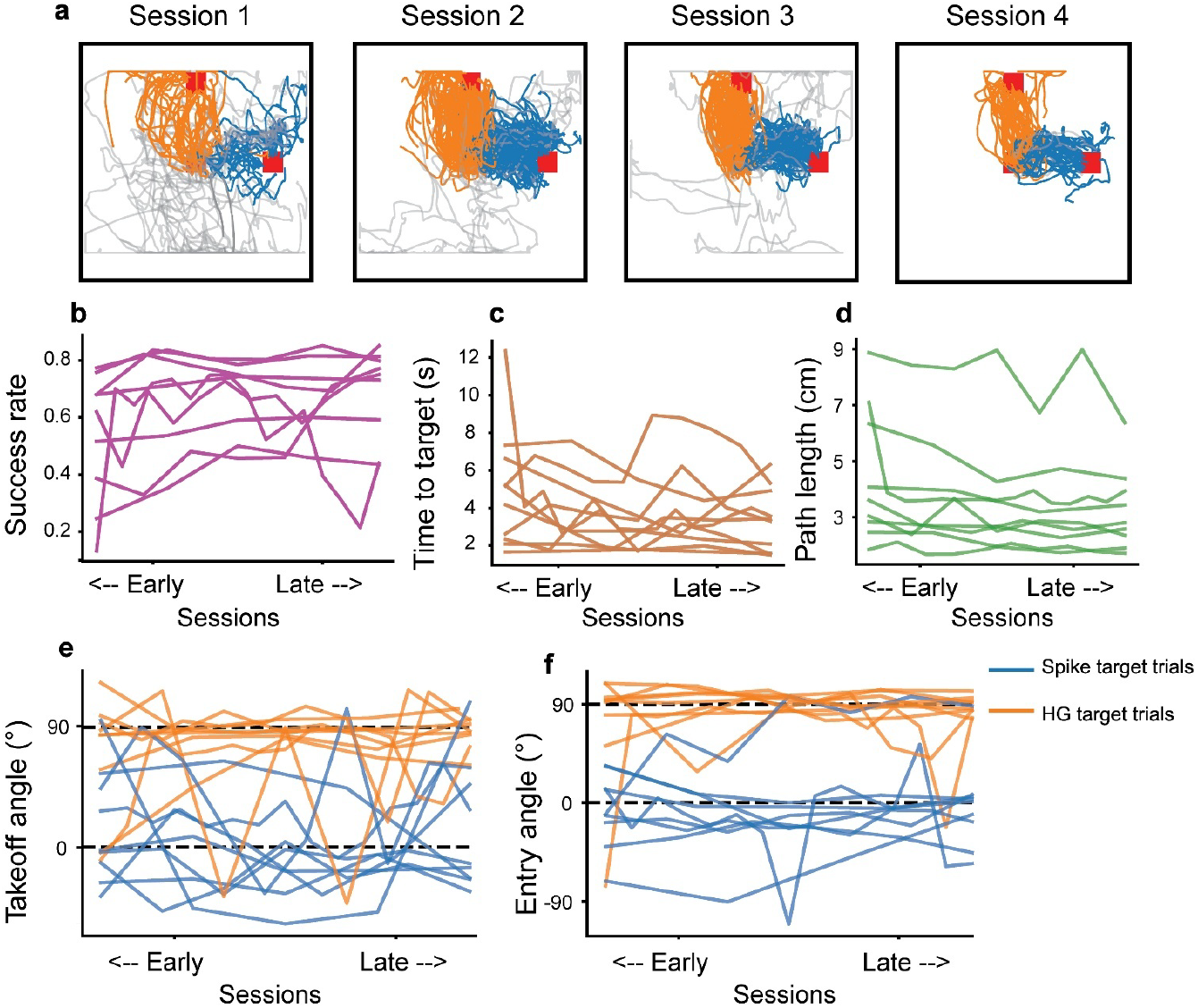
Performance on the ONF task. **a**, Cursor traces of the last 2 s of all trials in 4 sessions from one CE of monkey C. The red square in the center of the screen is the center target. The red square at the top represents the HG target (orange traces, individual HG target trials) while the red square on the right represents the spike target (blue traces, spike target trials). Grey traces indicated failed trials (see Methods). **b**, Success rate for all monkey-CE combinations of each session. Each line represents one combination; note that each combination was trained for a different number of sessions. **c**, Mean time to target of all trials for all monkey-CE combinations of each session. **d**, Average path length of all trials for all monkey-CE combinations of each session. **b-d**: Slopes were significantly different from zero (p<0.05, one sample t-test; see Methods). **e**,**f**, Average takeoff angles and entry angles of each monkey-CE combination over sessions (see Methods). Angles did not change significantly over sessions; rather, they were generally close to 0° or 90° (for spike or HG target trials, respectively) within the first session.

### HGA and spiking on the same electrode can be independently modulated during ONF control

To examine the independence of spiking and HGA, we plotted the normalized, trial-averaged spike rate and HGA in 50-ms bins for one example CE leading up to reward in monkey C during hand control and ONF control. During hand control (Fig. 4a), trial-averaged HGA and spike rate sometimes modulated together, but often did not. During ONF control (Fig. 4b), we observed a stark dissociation of the two signals. Specifically, during trials to the spike target (solid lines), spike rate increased while HGA did not; during trials to the HG target (dashed lines), the opposite occurred. To quantify the dissociation on a single-trial level, we computed the Pearson correlation coefficient (R) between single-trial spike rate and HGA from the last second of each trial (twenty 50-ms bins; see Methods). We compared this single-trial correlation during ONF control trials with the correlation during hand control trials for every monkey-CE combination (Fig. 4c) after the monkeys had become proficient at the task (defined as acquiring at least 40 successes per file). Note that HGA and spike rate were moderately, not highly, correlated even during hand control, either on the CE (0.36 ± 0.34, mean ± s.d.), or on other electrodes on the array (Extended Data Fig. 2a). Correlation values decreased significantly during ONF control in all CEs to near 0 (−0.037 ± 0.39, mean ± s.d.). This held true even in the electrodes with the highest hand-control correlation across different sessions, as well as during the first 1 s of trials (Extended Data Fig. 2b), and from the start of the first session in most cases (Extended Data Fig. 1c). Thus, the monkeys could independently modulate spikes and HGA recorded on the same electrode at a single trial level. As an additional control for the possibility that monkeys were just separating the closest 1-2 neurons from HGA on the CE, we lowered the spike threshold to 3 s.d. (in the “hash,” or multiunit spikes) for 2 CEs. While the single unit on the CE in this case was stable over sessions (days), the unsorted waveforms varied quite widely, implying they represented spiking of many local neurons (Extended Data Fig. 3a-d). Even in this case, the monkey could still independently modulate the two control signals (R= 0.03, 0.01; Extended Data Fig. 3e).

**Figure 4.**
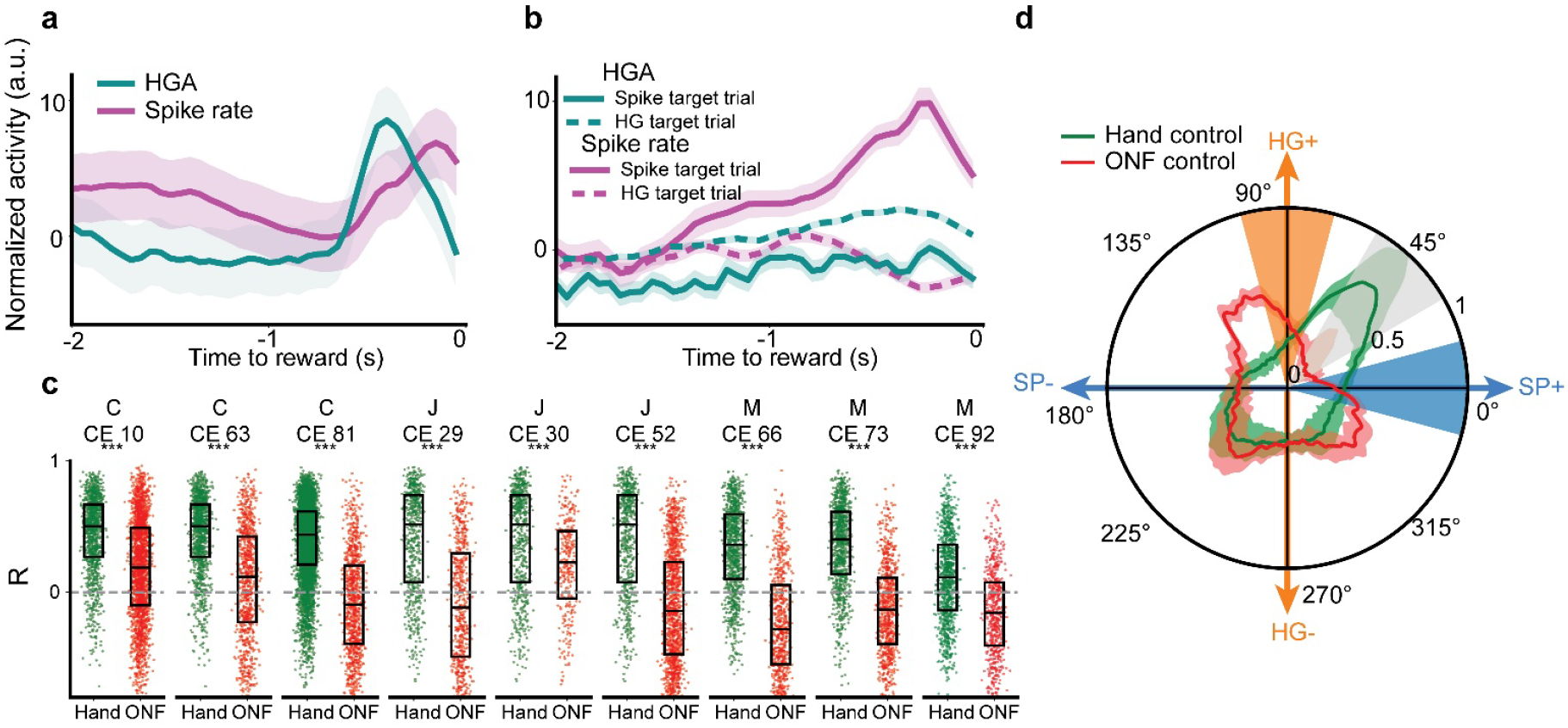
Independent modulation of spikes and HGA. **a**, Trial-averaged, normalized spike rate and HGA recorded on an example CE during the last 2 s of hand control when the monkey was acquiring the correct target. Shaded areas: standard error. **b**, Same as **a**, but for ONF control. Solid lines: spike target trials; Dashed lines: HG target trials; turquoise: HGA on the CE; magenta: spike rate on the CE; Shaded areas: standard error. **c**, Median (±IQR) correlation (R) between spike rates and HGA in hand control trials (green) and ONF control trials (red) for all monkey-CE combinations (letter-number). Each dot represents one trial. Correlations decreased significantly on all electrodes (***p<0.005, two-sided Wilcoxon rank sum test). **d**, Polar plot of the 2D distributions of mean (±s.d.) spike-HG angles of 20 time bins (1 s) preceding the reward time during all ONF (red) and hand control (green) from the proficient sessions over all monkey-CE combinations. Orange sector: HG+ state; blue sector: SP+ state. Gray sector: state at which both signals had high modulation.

Next, we sought to examine this independent modulation on the level of individual time bins. In the one second before reward, we defined the ‘spike-HG angle’ for each 50-ms bin as the angle formed by the vector sum of the mean HGA and spike rate on the CE for that bin (Fig. 4d, Extended Data Fig. 4). We then defined the SP+ state as times with increased spike rate and baseline level of HGA (spike-HG angle of 0°±15°) and the HG+ state as times with increased HGA and a baseline spike rate (spike-HG angle of 90°±15°). During ONF control across all monkey-CE combinations, a larger number of spike-HG angles were in the SP+ states (9.3%) and HG+ states (18.5%) than during hand control (SP+ states: 7.2%; HG+ states: 7.4%); i.e., there was more independent modulation during ONF control. Moreover, there were fewer bins with spike-HG angles indicating simultaneous increases in HGA and spike rate on the CE (45°±15°) during ONF control (3.6%) compared to during hand control (16.1%), showing decorrelation on short timescales. Taken together, these results consistently showed that HGA and local firing activity could be separated on the same electrode.

### HGA was correlated with distributed spiking activity but not due to spike leakage from neurons on neighboring electrodes

To further investigate the possibility that HGA is generated by leakage of spikes through volume conduction, we examined the relationship between HGA on the CE and spikes on all electrodes. First, we qualitatively examined the trial-averaged spike rate and HGA in each electrode in the time prior to reward for each type of trial (Fig. 5a-d, monkey C, CE 63). During spike target trials, spike rate increased most on the CE, with minimal changes in most other electrodes’ spiking (Fig. 5a) and without changes in HGA on any electrodes (Fig. 5b). During HG target trials, spike rate decreased on the CE and increased on multiple other electrodes (Fig. 5c), while HGA increased strongly on the CE and weakly on other electrodes (Fig. 5d). We then examined the spatial relationship across the array between mean spike rate and HGA over the last second of each trial (Fig. 5e,f; Extended Data Fig. 5). If HGA on the CE were due mainly to spike leakage, it would be reasonable to expect higher spike rates on the electrodes surrounding the CE on HG target trials, since spiking on these electrodes would contribute most to the increase of CE HGA when the CE spiking rate did not modulate. Instead, in HG target trials, we observed a widely distributed increase in spike rates across the array, with no bias toward electrodes neighboring the CE. These qualitative results suggest that HGA is not primarily driven by spike leakage from neurons on nearby electrodes. To quantify this observation, we tested whether HGA on the CE was significantly more correlated with weighted summed spiking activity near the CE than with summed spiking near randomly selected electrodes. We modeled (within the limits of Utah array sampling) the contribution of the nearby spiking activity to HGA on the CE by volume conduction. That is, we computed a sum over electrodes of trial-concatenated spike rates in each time bin weighted by the inverse of the square of the distance from the CE (see Methods). In this scenario, of course, the greatest contributor to HGA would be the spiking activity recorded on the CE itself. We found that this distance-weighted sum of spiking (DWSS) correlated minimally with trial-concatenated HGA on the CE (R=0.10 ± 0.12, N=9; Fig. 5g; Extended Data Fig. 6). This was not significantly larger than either (1) the correlation of CE HGA with the DWSS calculated using random electrodes, excluding the control electrode (one-sided Mann-Whitney U test, p>0.9), or (2) the correlation between HGA and the DWSS on the same random electrodes (one-sided Mann-Whitney U test, p>0.9; Fig. 5g; Extended Data Fig. 6; see Methods). Thus, nearby electrodes were not particularly predictive of the HGA on the CE.

**Figure 5.**
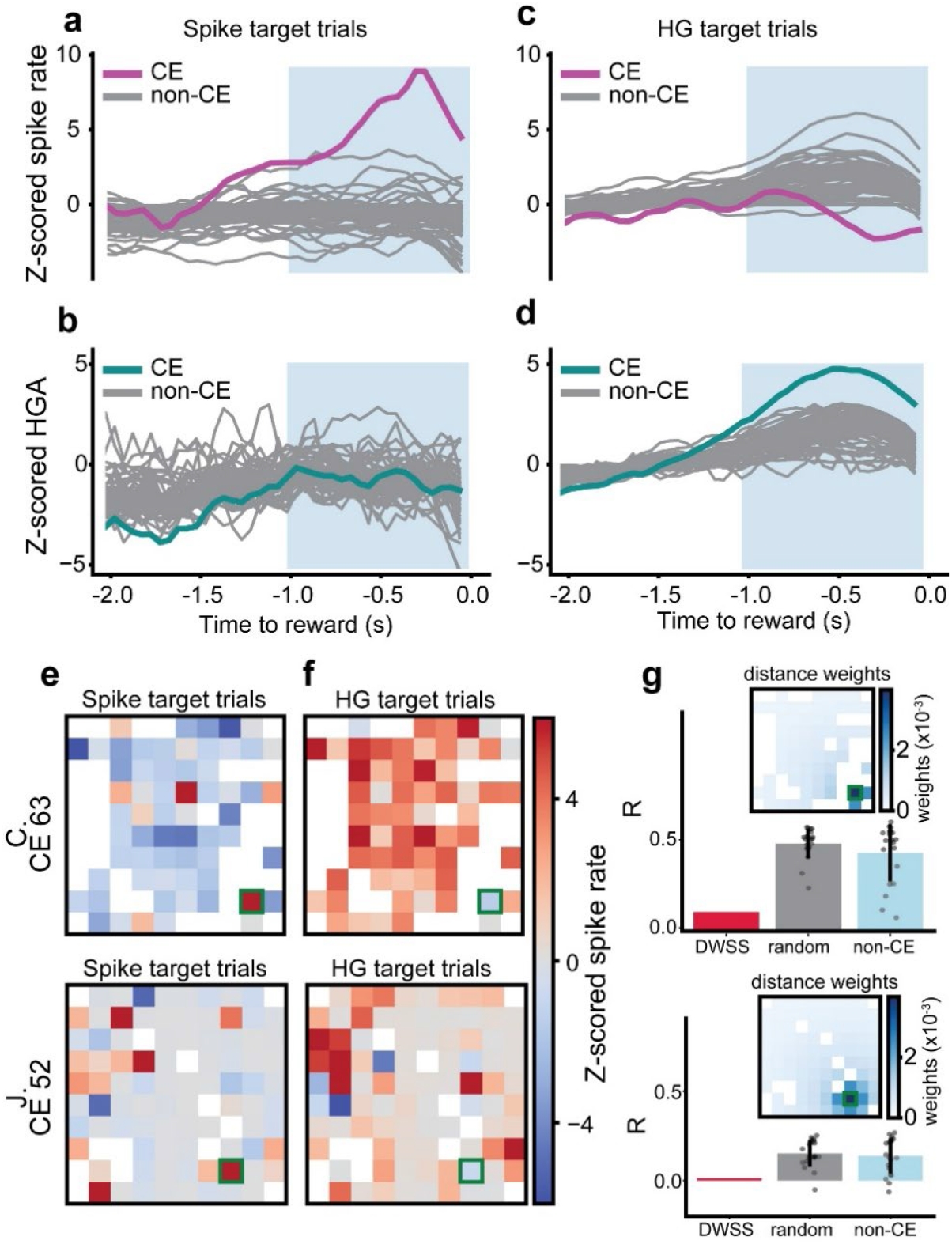
Neural spiking patterns across the array during ONF. **a, b**, Trial-averaged, z-scored spike rates (**a**, magenta) and HGA on the CE (**b**, green) and non-CEs (gray) during the last 2 s of all the spike target trials of ONF control, aligned to reward onset. **c, d** same as **a**,**b** for all the HG target trials of ONF control. Data **c, d** same as **a**,**b** for all the HG target trials of ONF control. Data of each electrode in **a**-**d** were z-scored by the population mean and standard deviation. Shaded area highlights the neural activities during the last second of the trial. **e, f** Trial-averaged, z-scored spike rates over the last 1 s of spike target trials (**e**) and HG target trials (**f**) of ONF control in each array of two example monkey-CE combinations. White squares denote shunted electrodes (see Methods). CEs are outlined in green. **g**, Correlation coefficient (R) between CE HGA and the CE distance-weighted sum of spikes (CE DWSS, red bar), the mean (±s.d.) R between CE HGA and DWSS of 20 randomly selected electrodes (rand DWSS, grey bar; each dot is one R with one electrode), as well as the mean (±s.d.) R between HGA and DWSS pairs from 20 randomly selected electrodes *excluding* the CE (non-CE, light blue bar). The R value between CE HGA and CE DWSS was not significantly larger than those between CE HGA and random DWSS, or than those between non-CE and DWSS (n.s., one-sample, one-sided Mann-Whitney U test). Insets: the weights used to compute the CE DWSS. **e**-**g**, Top row: monkey C, CE 63; Bottom row: monkey J, CE 52.

**Figure 6.**
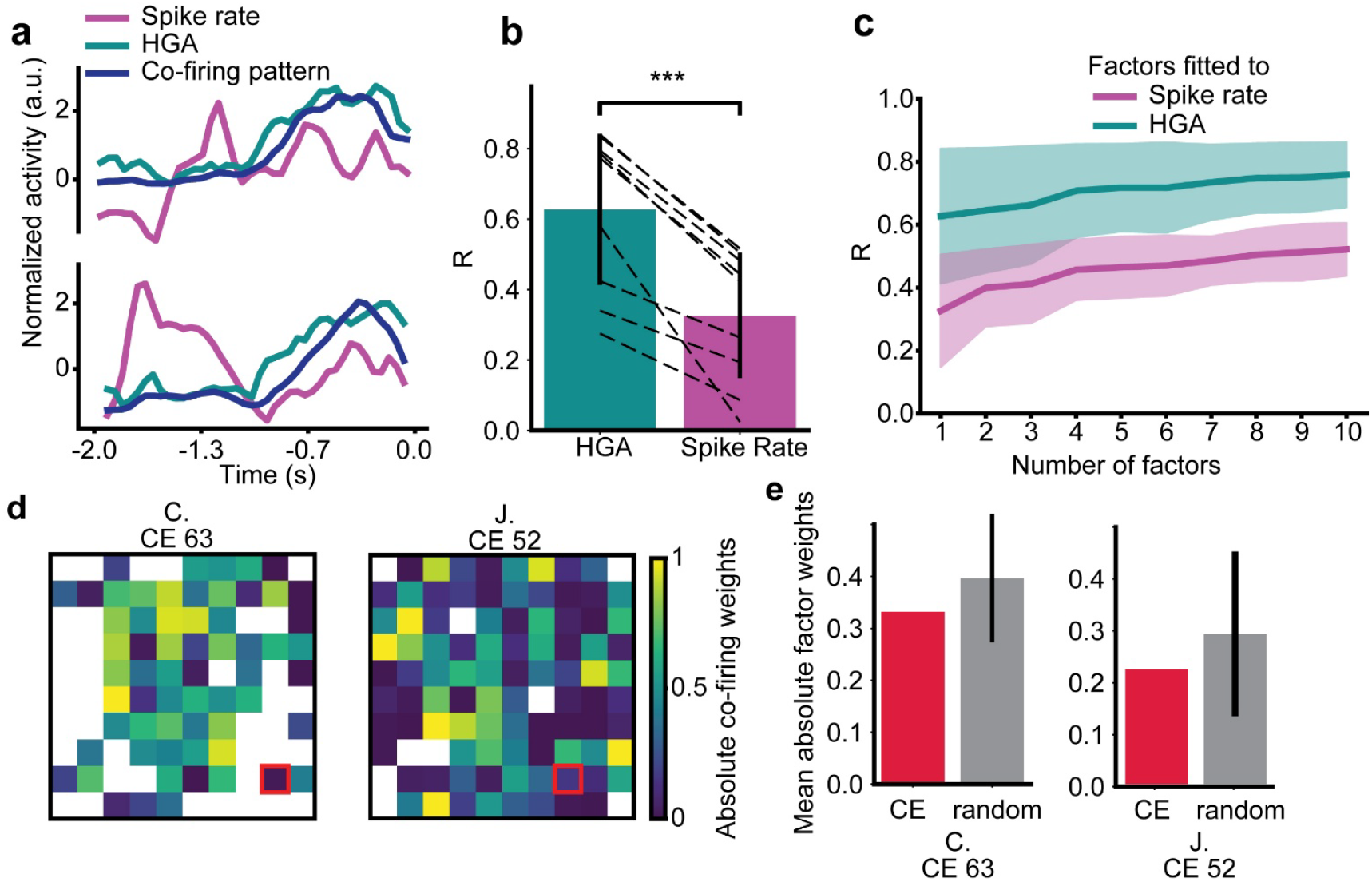
HGA on the CE correlated more with co-firing than the CE spike rate did. **a**, Single trial examples comparing the first co-firing pattern (dark blue) to HGA (turquoise) and spike rate (magenta) on the CE (time relative to reward onset). **b**, Correlation (R) of multivariate linear regression (MLR) fit of the first co-firing pattern to spike rate (magenta), or to HGA (turquoise), averaged over all the CEs (dashed lines, individual CEs, N=9; bars, mean ± s.d., ***p=0.004, two-sided Wilcoxon signed rank test). **c**, R of MLR fit using an increasing number of factors to either spike rate (magenta) or HGA (magenta). We fitted a factor analysis model for each number of factors separately. Thick trace and colored shades: mean, s.d. over CEs. **d**, Absolute values of the co-firing weights of two example monkey-CEs. CEs outlined in red. **e**, Mean values of the absolute co-firing weights on the electrodes adjacent to the CE (red), as well as the mean (±s.d.) of average value of the absolute co-firing weights on the electrodes adjacent to 20 randomly selected electrodes (gray) of two example monkeys. These R values were not significantly different (p>0.4, one-sided permutation test).

### HGA correlated with neuronal population co-firing patterns across the array

In the previous section, we observed that spike rates increased on many non-CEs in HG trials (Fig. 5b,d), suggesting a possible connection between mesoscale population-level spiking and HGA on the CE. To explore the relationship, we applied factor analysis to the spike rates of all electrodes after the monkeys had become proficient at the task to extract the patterns of greatest neuronal covariation. We used a 1-factor model to extract the common factor that best explained the spiking activity over many days. We call this factor the “co-firing pattern”^8^, which is a low-resolution measure of synchrony. We first examined the relationship between the co-firing pattern and spiking and HGA on the CE for randomly selected single trials (Fig. 6a). The co-firing pattern was highly correlated with CE HGA but uncorrelated with CE spiking. Indeed, this relationship held across all monkey-electrode combinations – regression models between co-firing and HGA (R=0.6 ± 0.13, mean±s.d.; Fig. 6b) had significantly better fits than between co-firing and spike rate (R=0.3 ± 0.11). The relationship between co-firing and HGA also held in individual sessions (Extended Data Fig. 7a) and became even more pronounced when co-firing was described by factor analysis models with an increasing number of factors (Fig. 6c). That is, the HGA was highly correlated with the low-dimensional activity of the neuronal ensemble.

**Figure 7.**
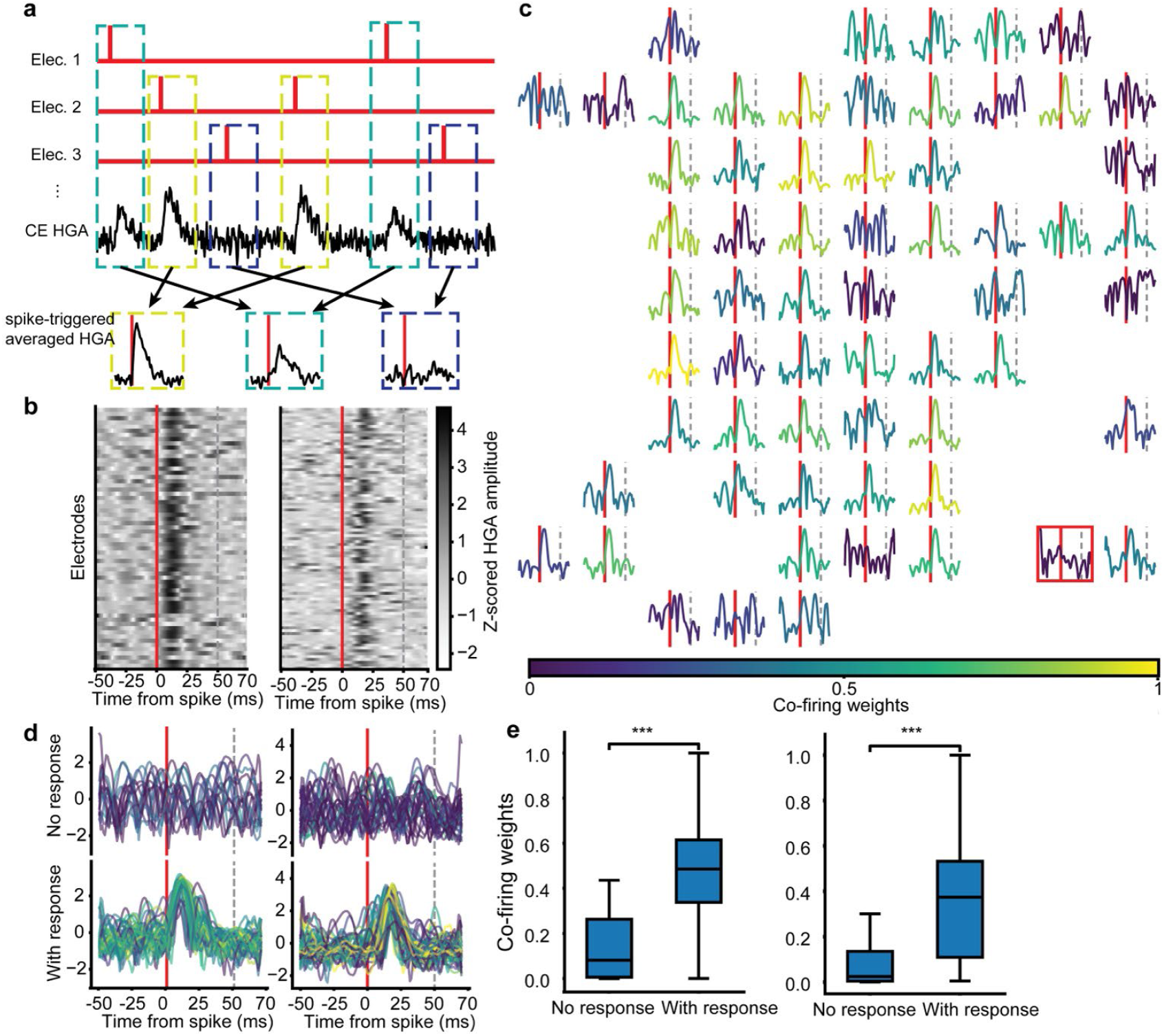
Spikes contributing most to co-firing also contributed most to HGA. **a**, An illustration demonstrating the calculation of mean CE HGA triggered by non-CE spikes. For all spikes on a given electrode, we averaged the corresponding CE HGA segments from −50 to 70 ms relative to the spike times to obtain the spike-triggered HGA for that electrode. **b**, Spike-triggered averaged HGA of the CE (two examples shown) showed a consistent activity pattern when aligned to the spike times of other electrodes. It peaked about 12.5 ms after spike time. **c**, An example of spike-triggered average HGA on the CE (red square) of monkey C, CE 63, plotted on each electrode from which the spike time was used to align the HGA. Red vertical line represents the spike time; the gray dashed line represents 50 ms after the spike time. HGA traces are color-coded by the absolute values of the co-firing weights from **Fig. 6d. d**, K-means clustering results using 2 clusters on the STA HGA traces, color-coded by the absolute co-firing weights. Cluster with apparent peak: “With response”; without apparent peak: “No response”.) **e**, A comparison of the absolute co-firing weights of the two clusters (***p<0.005, two-sided Wilcoxon signed-rank test). **b**,**d**,**e**, Left: monkey C, CE 63; Right: monkey J, CE 52.

We next investigated the spatial patterns of the electrode contributions to the co-firing pattern during ONF control. We observed that the electrodes with high co-firing pattern weights were spread broadly across the array (Fig. 6d). This was true in all three monkeys, with variation in the numbers of highly weighted electrodes across monkeys (Extended Data Fig. 7b). Notably, there was not a high concentration of weights in electrodes closest to the CE (Fig. 6e; Extended Data Fig. 7c), consistent with our prior DWSS results.

To further examine how population spiking activity across the array related to CE HGA and spike rate, we constructed neural subspaces unique to the times at which the monkey was modulating only HGA or only spike rates on the CE after the monkeys had become proficient at the task (i.e., the HG+ and SP+ states, Fig. 4d). A neural space can be defined as an N-dimensional state space with each electrode’s spike rate mapped to one of the N dimensions. We used contrastive principal component analysis (cPCA, see Methods) to find lower-dimensional neural subspaces of the N-dimensional neural space that maximized variance in one spike-HG angle (e.g. HG+) and minimized it in the other (e.g. SP+). We projected the trial-concatenated spike rate data of all successful ONF trials into a SP+ cPC subspace and a HG+ cPC subspace and correlated those two projections with the HGA and spike rate on the CE from the same trials. As we expected, across all monkey-CE combinations, activity in the HG+ subspace correlated with CE HGA, while activity in the SP+ subspace correlated with the CE spike rate (Extended Data Fig. 8a). This relationship also held for individual sessions (Extended Data Fig. 8b). We also observed that the highly weighted electrodes for the first HG+ cPC were distributed across the array with no significant concentration around the CE (Extended Data Fig. 8c,d; one-sided permutation test, p>0.05). If the spiking activity during HG+ states was more correlated with CE HGA, we would also expect more synchronous (co-firing) activity occupying the HG+ neural subspace, leading to lower dimensionality of the HG+ subspace. We examined the cumulative variance accounted for of the neural subspace of HG+ and SP+, defined by singular value decomposition, and observed that the HG+ subspace was significantly lower dimensional than the SP+ subspace (p=0.02, permutation test on mean difference curve; Extended Data Fig. 8e). Overall, these results supported the prior findings that population co-firing patterns have a higher correlation with HGA (Fig. 6b,c), and that the population responsible for these patterns need not be concentrated around the CE (Fig. 6d,e).

### Control electrode HGA peaked shortly after spikes from the electrodes that contributed most to co-firing

The above analyses linked the CE HGA with synchronous population activity, or co-firing. How does synchronous population firing generate CE HGA? One possibility is the second hypothesis above (Fig. 1): synchronous spikes from broadly distributed neurons arrive at dendrites and/or somas of neurons near the CE and elicit postsynaptic potentials (PSPs) that summate. In this case, we would predict an increase of CE HGA following the spikes recorded on other electrodes. Further, spikes on the electrodes that contributed more to the co-firing pattern (which correlated with HGA) should contribute more to the increase of CE HGA. To investigate these relationships, we computed the mean CE HGA triggered by the spikes on each electrode during ONF control, or spike-triggered average HGA (Fig. 7a; Methods). We observed an increase of the CE HGA triggered by spikes on most of the other electrodes (Fig. 7b; some electrodes’ spikes did not elicit an increase in HGA). This increase of CE HGA was observed across sessions as well (Extended Data Fig. 9). The CE HGA peaked at a mean lag of 10.2 (95% CI: 5.9 - 14.5) ms and 10.9 (95% CI: 6.0 - 15.8) ms after the spike time for monkeys C and J, respectively (Fig. 7b). For monkey M, the mean peak delay was 16.5 (95% CI: 13.5 - 19.5) ms. This range of delays agrees with estimated timing of the peak of postsynaptic potentials relative to presynaptic spike times in a prior study^47,45^. Furthermore, spikes on electrodes with higher co-firing weights (Fig. 6d) produced a larger peak in the spike-triggered HGA average (Fig. 7c). To characterize this observation, we used k-means clustering to separate the normalized spike-triggered HGA traces into 2 clusters based on their shape. This unsupervised algorithm produced two groups of average spike-triggered HGA traces with distinct shapes (Fig. 7d). One cluster (“With response”) showed a clear increase in HGA following the spikes, while the other had no clear relationship between spiking and HGA (“No response”). The co-firing weights of the corresponding electrodes in cluster “With response” were significantly higher than the weights in cluster “No response” (two-sided Wilcoxon rank sum test. Monkey C, CE 63, p=0.003; monkey J, CE 52, p=0.001, Fig. 7e). These effects were observed in all monkeys (Fig. 7d,e; Extended Data Fig. 9). Thus, the neurons that contributed most to the network co-firing pattern also contributed most to the spike-triggered HGA average, which peaked 10 - 15 ms after the spike times. Combined with the fact that the co-firing pattern correlated highly with HGA, these results support the hypothesis that synchronized firing activity from the neuronal population arrived at, and elicited synchronized PSPs from, dendrites and somas close to the CE.

### Independent modulation of spikes and HGA was an intrinsic property of the neural population

The results in Fig. 6 indicated the existence of unique subspaces for CE spike rate and HGA. This enabled us to further investigate how these signals were dissociated in our ONF BMI task. The independent control of the two signals was reflected in the close-to-orthogonal relationship between the leading subspace principal angles of SP+ cPCs and HG+ cPCs (Extended Data Fig. 10a). We questioned whether the monkeys had to learn completely new ensemble firing patterns to modulate CE HGA or spike rates in isolation. If this were true, the SP+ and HG+ cPCs would explain close to no variance of the hand control data that span the “intrinsic manifold.” To test this, we projected the population spiking activity during hand control into the HG+ and SP+ cPC subspaces and computed the variance accounted for in the original hand control data by these two unique subspaces (Extended Data Fig. 10b,c; Methods). Over all nine control electrodes, the top 10 HG+ cPCs and SP+ cPCs explained 16.7 ± 2.7% and 9.7 ± 1.3%, respectively, of hand control data variance. In comparison, the top 10 HG+ cPCs and SP+ cPCs computed from hand control data explained 26.3 ± 7.0% and 15.9 ± 4.3% of hand control data variance. Thus, the cPCs used for ONF control account for over half of the hand-control-only cPCs comparison. (For reference, the top 10 PCs of hand control data (intrinsic manifold) explained 60.9 ± 4.0% of its own variance.) These results suggested the ability to independently modulate HGA and spiking lay partially within the intrinsic neural manifold, which implies that the findings about the source of HGA were not unique to ONF. That is, HGA and spike sources are independent, which fits with the observation that monkeys became proficient in the task quite quickly.

## DISCUSSION

High gamma activity has often been characterized as a proxy for spiking near the recording electrode^1,2,20–22,24–28^. Here, we demonstrated causal evidence that subjects could use a novel BMI to actively modulate spiking activity independently from HGA on the same electrode. Subjects were able to quickly and consistently decouple spiking and HGA, even with a very low spike threshold. Further, we demonstrated that HGA was correlated with synchronized population spiking activity (co-firing) that was distributed widely across the array, rather than a distance-weighted sum of spiking. Spike-triggered HGA on the CE peaked about 12.5 ms after spikes occurred on other electrodes and was highest for spikes from electrodes contributing most to the co-firing pattern. Finally, we identified two orthogonal neuronal subspaces that were associated uniquely with HGA modulation and spiking modulation on the control electrode, respectively. These subspaces overlapped with the neuronal manifold active during arm movement. Together with the rapid ability to decouple spiking from HGA, this suggests that this decoupling was intrinsic, rather than learned.

Multiple influential studies contributed to a prevalent view of HGA as an index of summed spiking. Our results in fact largely agree with the findings that led to this view. First, the finding that co-firing of distributed neurons correlates highly with HGA agrees with prior findings of high correlations of HGA with spiking that was averaged over broadly spaced electrodes. By contrast, those seminal studies showed, as we (Extended Data Fig. 2) and others^16,38^ also found, only modest correlations between HGA and spikes on single electrodes. While experimental^31,35^ and computational^23^ modelling studies showed that spike leakage can contribute to HG band, the proportion contributed was modest^37^.

Our results also concur with findings from multiple computational models^2,46,47^ and experimental studies^48–52^ suggesting that HGA is generated by summed postsynaptic potentials^23,53^ and possibly dendritic calcium spikes^33^. HGA can increase regardless of local spiking changes^54^ and retains decodable information in the absence of recorded spiking activity on the same electrode^9,50,55–57^. Most convincingly, blocking NMDA-mediated postsynaptic activity reduced HGA, but not spiking activity^39^. Here, we saw that spiking recorded on the CE, which should contribute the most to leakage, could be voluntarily decoupled from HGA, even from the highest spectral portion of HGA (200-300 Hz). Moreover, monkeys could decorrelate spikes from HGA even with a spike detection threshold in the hash, resulting in many neurons contributing to the spiking signal (Extended Data Fig. 3). HGA strongly correlated with co-firing across the broader network, yet our DWSS analysis showed that this was not due to leakage from broadly spaced neurons. Our spike-triggered average HGA results showed that the CE HGA peaked from 6-20 ms after spikes on other (non-CE) electrodes. Importantly, this agrees with PSP timing relative to presynaptic spikes from prior work^45^. Our findings thus agree with and extend this prior evidence.

Experimental studies and computational models have suggested that synchronous inputs greatly increase the amplitude of intracortical LFPs^53,58^ and HGA in ECoG^59^. A more nuanced view showed that the latent dynamics of the broader neuronal population during reaching correlated with HGA on individual electrodes after alignment procedures^38^. Here, we found that both single factor co-firing and the cPCs for HG+ states correlated highly with HGA without the need for subspace alignment. Furthermore, the co-firing pattern we saw involved neurons that were distributed across the array, while the contribution of the neighboring electrodes was not particularly high. Our results thus corroborate and extend these studies, showing that mesoscale synchrony may be important in generating local HGA.

Overall, our findings help to reconcile prior evidence of the importance of PSPs, synchronous inputs, and spike leakage to HGA. The hypothesis that synchronous PSPs are the predominant source of input, with leakage being a lesser contributor, is the most parsimonious way to harmonize our findings with prior observations. In this hypothesis, the high correlations of spatially averaged spikes with HGA are likely due to both signals being driven by the same underlying synaptic events: PSPs generate HGA and increase the probability of nearby action potentials. A hypothesis in which HGA arises primarily from spectral leakage of local spikes has more difficulty explaining the active and rapid dissociation between HGA and spikes that we observed on multiple different CEs in different monkeys, even with a very low spike threshold, the low correlations of HGA and local spikes in most cases even during hand control, and the strong relationship between HGA and spikes recorded millimeters away. PSPs summate to higher levels when the inputs generating them are more highly synchronized. Their much longer duration than spikes also makes summation easier, as spikes would have to synchronize within 1 ms to summate. Thus, a PSP source of HGA would predict synchronous firing, but not spatial concentration, of input neurons. We found synchronous firing across the array that correlated highly with HGA and contributes highly to spike-triggered HGA at a lag consistent with synaptic conduction time. Therefore, our observations are most simply explained by the predominant source of HGA being summed PSPs in somas and/or dendrites near the recording electrode generated by synchronous spiking of mesoscale-distributed presynaptic neurons.

This study has some limitations. While the interspike interval histograms and highly variable waveforms in Extended Data Fig. 3 suggest that our spike control signal represented many neurons, we cannot precisely quantify that number of neurons. Additionally, the Utah array has limited spatial resolution. Thus, we cannot completely rule out the possibility that a considerable source of HGA is unobserved neurons, either within or outside the plane of the array. Still, for this to happen, those neurons would need to be near enough to leak into HGA band recorded on the CE but not near enough for their spikes to exceed the detection threshold (i.e., more than 140-200 μm away^60,61^). Future work with higher density electrodes and broader coverage could help investigate the possibility of leakage further and whether neurons even farther away than 4 mm contribute to local HGA.

In summary, the generation of HGA is likely a complex phenomenon arising out of interactions of various types of neurons embedded in a cortical network in a spatially distributed manner. This characteristic potentially provides HGA with several benefits for both fundamental and applied research. Because HGA appears to be predominantly generated by summed PSPs produced by inputs from a distributed neural population, it is more robust than spike recordings to electrode movement^62,63^ and neuronal death or migration^64,65^, which gives HGA greater longevity than spike recordings with current electrode technologies. In addition, this distributed contribution to local HGA reduces sensitivity to variability in individual neuron firing patterns, which makes it more stable over time than spikes^9,66^ without additional procedures like subspace alignment^38,67^. These factors are attractive for BMI applications. In addition, the fact that HGA represents synchronous, widely distributed inputs could be useful in studies of integration of inputs from other brain areas to local neurons, which is important in sensory integration, attention, language, and decision making^3,4,11,13–17^. On the other hand, this fact may limit conclusions that can be made using HGA about activity in individual neurons in the neighborhood. HGA can also serve as a parsimonious marker for mesoscale synchronous firing, which has been proposed to be involved in a wide range of brain functions, including attention and binding^14,68–70^. While we only examined M1, it seems plausible that these findings would apply to many other neocortical areas that have similar cytoarchitectures. Thus, these findings have ramifications for a wide range of neuroscience and neuroprosthetic investigations.

## METHODS

All procedures were performed under protocols approved by the Northwestern University Institutional Animal Care and Use Committee. Three adult rhesus macaque monkeys were trained in the experimental procedures described below.

### Surgery

In each monkey, we surgically implanted a 96-electrode silicon electrode array (Blackrock Microsystems) into the primary motor cortex (M1) contralateral to the arm used to control the cursor. During surgery, monkeys were anesthetized with isoflurane (2–3%). Anesthetic depth was assessed at all times by monitoring jaw muscle tone and vital signs. A craniotomy was performed above M1, and the dura was incised and reflected. Electrode arrays were implanted in the proximal arm area, as determined by referencing cortical landmarks and by intraoperative electrical stimulation with a silver ball electrode (2–5 mA, 200 μs pulses at 60 Hz). During stimulation, a reduced level of isoflurane (< 0.5 MAC) was supplemented with intravenous remifentanil (0.15 -- 0.30 μg/kg/min) to reduce suppression of cortical excitability. The electrode array was positioned on the crown of the right precentral gyrus, approximately in line with the superior ramus (medial edge) of the arcuate sulcus. Electrode shank length was 1.5 mm. The preamplifier was grounded to the Cereport pedestal and referenced to two platinum wires with 3-mm exposed wire length placed under the dura. A piece of artificial pericardium (Preclude, Gore Medical) was applied above the array, and the dura closed using 4.0 suture (Nurolon, Ethicon). Another piece of pericardium was applied over the dura, and the craniotomy partially filled with two-part silicone (Kwik-Cast, World Precision Instruments). The craniotomy was then closed, and the skin was closed. All monkeys were given postoperative analgesics buprenorphine and meloxicam for two and four days, respectively.

### Neural signal recording

All neural and behavioral data were recorded using a 96-channel Multiple Acquisition Processor (Plexon, Inc). LFP signals were sampled at 1 kHz and band-pass filtered from 200 to 300 Hz. We deliberately chose this band to be as conservative as possible, as prior studies suggested that components of LFPs above 150 Hz were most contaminated by spike leakage^31,34,35^. We computed the spectral power of each electrode from the band-passed LFP by applying a 256-point Hanning window (overlapped by 206 points) and calculating the squared amplitude (power) of the windowed signal’s discrete Fourier transform, yielding 50-ms binned HGA. Multiunit spikes (threshold crossings) were high-pass filtered at 300 Hz and sampled at 40 kHz. The signals were further downsampled to 1 kHz.

The threshold on each electrode was set at 4.5 s.d. from the root-mean-square baseline activity on the electrode. Note that we also ran a subset of experiments using a lower threshold of 3 s.d. (i.e., in the “hash”, to reduce the possibility that we were missing some smaller-amplitude spikes near the electrode) and obtained similar results - monkeys could still decorrelate the signals. We defined spikes as unsorted threshold crossings on each electrode. To identify shunted electrodes, we calculated the pairwise R of the spike rates, binned at 1 ms, between all electrode pairs and removed any electrode that had a pairwise R higher than 0.3 with any other electrodes. For all the analyses except the spike triggered HGA analysis, we used 50-ms bins for both HGA and spike rates. The above procedures were performed for each file separately. In each file, we also removed electrodes on which the spike rates averaged less than 5 spikes/s. The available electrodes were used for further analysis after these shunted and low-spike rate electrodes were removed.

### Hand control task

Three adult rhesus macaques, two male and one female (monkeys C, J, and M), used a two-link manipulandum to move a cursor (1-cm diameter circle) within a rectangular planar workspace (20 cm × 20 cm). The task was a four-target, center-out task, with 2-cm square outer targets spaced at 90° intervals around a circle of radius 10 cm. Each trial began with a target at the center of the circle. After a random hold time of 0.5 -- 0.6 s within the center target, a randomly selected outer target was illuminated and the center target disappeared, signaling the monkey to start a reach. The monkey needed to reach the outer target within 1.5 s and hold for a random amount of time between 0.2 s and 0.4 s to obtain a liquid reward. The task was performed in 10-minute-long files.

### Orthogonal Neurofeedback (ONF) task

To maximize the ability for monkeys to control the cursor, we first selected electrodes on which spiking correlated highly with movement during the reach task. First, we computed the absolute value of the Spearman correlation coefficient (|R|) between binned spikes and HGA on each electrode and the velocity in the X and Y directions, respectively, for a total of four correlations (spikes-X, spikes-Y, high gamma-X, high gamma-Y) over a 10-minute period. Next, we added the |R| corresponding to X and Y directions for each signal (i.e. spikes-X + high gamma-Y; spikes-Y + high gamma-X) giving us two |R|s corresponding to the two possible arrangements of the signals for control. We then selected the electrodes with highest cumulative |R|(control electrodes, CEs) as inputs to use for the ONF task. Later, after monkeys learned to perform ONF control, we selected electrodes that had higher spike-HGA correlations (see below) during hand control.

ONF control was implemented by mapping linear filter outputs of the spike rate and high gamma signals to the X and Y components of cursor velocity, respectively. For either type of signal, we used a linear filter with a set of positive, identical weights set manually on the prior 5 time bins (50 ms each) of binned spike rates. A bias term was added to the filter output to allow the monkey to reach all parts of the workspace. A dynamic moving mean (averaged over the last 1 min of values) was subtracted from both HGA and spikes before applying filter weights to center the signals to avoid drift and allow for negative cursor velocity when necessary. Velocity was then integrated to obtain the updated cursor position. Thus, the cursor would have zero velocity when both signals were at their mean activity levels. Both velocity and cursor position signals were computed at 20 Hz resolution.

Initial training involved performing a 1D BMI task in each of the spike and HG directions. In this case, one 4-cm square target was placed along either the X or Y axis only, respectively, and the cursor moved directly toward or away from the target. The trials were the same structure as for hand control, except that the hold time was 0.1 s in both center and outer targets, with a maximum trial time of 15 s for a reward. Trials of which the cursor never reach the target within 15 s or fail to stay in the target for less than 0.1 s were considered as failed trials. A typical session (day) lasted approximately 60-90 minutes (6-9 files), with 10 minutes (1 file) performing hand control and 50-80 minutes (5-8 files) on BMI control.

Once the number of successes within a file exceeded 40 (typically after 3-7 days), the monkeys were switched to ONF (2D BMI) control. In this case, both filters were applied to their respective signals, and the outputs were mapped to X and Y velocity simultaneously –i.e., the cursor moved as a vector sum of filtered spikes and HGA. Targets in this task were the same as in the 1D BMI task, except that they were randomly placed on the X or Y axis in each trial. The rest of the task parameters (hold time, max trial time) were the same as in the 1D task. Because the cursor moved as a vector sum, ONF required the monkeys to increase either spike rate or HGA, while maintaining the other constant, to successfully reach the targets. Target sizes were gradually reduced to operantly condition the monkeys to learn to independently modulate spike rate and HGA. We performed ONF on 10 control electrodes across 3 monkeys; one was excluded due to noise affecting the spike recordings.

### Computing correlation between spike rate and HGA

To compare the correlation between spike and HGA for each control type (ONF control/hand control), we calculated Pearson correlation coefficient between single-trial spike rate and HGA of the last second (20 time bins) preceding reward during either ONF control or hand control of every monkey-CE combination. We summarized the distribution of R values for hand control (Fig. 4c, green) and ONF control (Fig. 4c, red) for every monkey-CE combination.

### ONF performance metrics

We measured performance in the ONF task using success rate, time to target, path length, takeoff angle and entry angle. We defined success rate as the number of successful trials divided by the total number of trials in each session. We defined time to target as the mean time from the outer target appearance to reward over all trials in each session. We defined path length as the mean cursor path of all the trials in each session. We defined takeoff angle as the mode of the distribution of angles formed by the vector of the change in cursor position from 0 to 500 ms after the go cue for all trials in each session. Similarly, we defined entry angle as the mode of the distribution of angles formed by the vector of the change in cursor position in the last 500 ms of the trial for all trials in each session.

Takeoff and entry angles were further grouped by the target type (spike target vs HG target). To examine if there were significant changes over sessions and monkey-CE combinations in success rate, path length, or time to target, we computed the slopes of the linear fits of the curves of these metrics of every CE across sessions and tested whether the slopes differed from 0 using one-sample t-tests. Similarly, to examine whether the takeoff and entry angles converged to 0° (for spike-target trials) or to 90° (for HG-target trials) over sessions across all monkey-CE combinations, we used one-sample t-test to test if the absolute values of the slopes of the fits to these curves were smaller than 0.

### Distance-weighted sum of spikes

To examine whether spike leakage could explain the HGA during ONF, we computed the distance-weighted sum of spikes. This assumed the contribution of spike-related activity to HGA on the CE was inversely proportional to the square of the Euclidean distance between the neuron spiking and the CE^71^, and assumed an isotropic medium, which is reasonable within a single cortical layer, which is the case for a Utah array. We calculated the distance-weighted sum of these spike rates from all available electrodes and treated this sum as the contribution of spike leakage to the CE HGA. We believed this was a reasonable approximation, even with the somewhat sparse sampling of a Utah array, since the listening radius for spikes on an electrode has been estimated at 140-200 μm, so most of the neurons within the plane of the array would be recorded by electrodes spaced apart by 400 μm. This distance-weighted sum of spikes (DWSS) of CE *c* was defined as

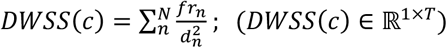

where *fr*_*n*_ ∈ ℝ^1×*T*^ was the concatenated spike rate of the last 2 s (40 bins) of all successful ONF control trials recorded on electrode *n, T* was the number of successful trials × 40 bins, *d*_*n*_ was the Euclidean distance between the electrode n and CE, and N was the number of all available electrodes. When n = c, we assumed that the spikes recorded on the CE were from neurons toward the outer part of the listening sphere (*d*_*c*_ = 140 μ*m*, a conservative estimate)^61^ from the recording site, minimizing the impact of the CE spiking on the DWSS. The correlation coefficient (R) between DWSS and HGA of the last 2 s (40 bins) of all concatenated trials on CE *c* could be written as

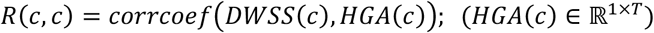

To understand the significance of the R value, we compared it to two controls. First, we computed the DWSS based on spike contamination for 20 randomly chosen electrodes *c*_*random*_. For any random electrode, the correlation coefficient between DWSS for the random electrode and HGA on the control electrode can be written as

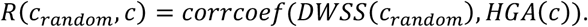

Then, we also calculated the correlation coefficient between DWSS and HGA on that same *c*_*random*_:

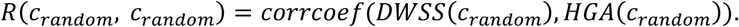

If the value of *R*(*c, c*) were significantly larger than the distribution of *R*(*c*_*random*_., *c*), or the value of *R*(*c, c*) were not significantly larger than *R*(*c*_*random*_, *c*_*random*_) (compared using Mann-Whitney U test), it would indicate that the spiking activity on the electrodes nearby the CE contributed significantly more to the HGA recorded on the CE than spiking activity surrounding a random electrode (Fig. 4g). That is, it would suggest that spike leakage was a significant contributor to HGA.

### Factor analysis

To extract the co-firing patterns for every monkey-CE combination, we built a factor analysis model from the concatenated spike population activity (***X*** ∈ ℝ^*N*×*T*^) recorded on all available electrodes of all successful ONF control trials, where *N* was the number of electrodes and *T* was the number of time bins of all concatenated trials. A factor analysis model sought to explain the correlations of spiking activities among electrodes by assuming they were influenced by common underlying factors (co-firing). Given the number of factors *n*, ***X*** was modeled as ***X*** = ***L*** ∗ ***F*** + ***R***, where ***L*** ∈ ℝ^*N*×*n*^ was the co-firing weights, ***F*** ∈ ℝ^*n*×*T*^ was the co-firing component, and ***R*** ∈ ℝ^*N*×*T*^ represented the noise independent to each neuron. The goal was to estimate ***F*** and ***L*** such that the variance-covariance structure of ***X*** was well explained by the common factors, thereby reducing the dimensionality of ***X*** while preserving its essential patterns. In Fig. 6a,b,d and Fig. 8, we set *n*=1 to extract the most common co-firing component and its corresponding weights. In Fig. 6c, we set *n*=1-10 to investigate how the number of factors affected the correlation between the co-firing components and CE spike/HGA.

### Contrastive principal component analysis (cPCA)

We used cPCA to find subspaces in which the monkeys were modulating only HGA or only spikes independently. cPCA found a subspace that maximized variance in the HG+ data (foreground data) and simultaneously minimized variance in the SP+ data (background data)^72^. The same process was used to find SP+-unique components (SP+ cPCs) but using SP+ data as foreground data and HG+ data as background data. cPCs were the eigenvectors of eigenvalue decomposition result on the matrix (***C***_*foreground*_ − ***αC***_*background*_) ∈ ℝ^*N*×*N*^, where ***C***_*foreground*_ ∈ ℝ^*N*×*N*^ and ***C***_*background*_ ∈ ℝ^*N*×*N*^was the covariance matrix of the foreground data and background data, respectively. cPCA has one hyperparameter *α*, which balances the importance of maximizing variance in the foreground data versus minimizing variance in the background data. We found that an *α* value of 10 was sufficient to separate the two conditions. We chose the top 10 cPCs ranked by their row variances for all further analysis. Projections into cPC unique space were calculated by projecting the entire trial-concatenated ONF control data to the cPCs.

### Subspace angle analysis

The principal angles between two subspaces provided a measure of the “alignment” of those subspaces in a high-dimensional space. These angles were defined recursively, capturing the smallest angles between vectors in the two subspaces. To compute the angles between two m-dimensional subspaces A and B embedded in a high dimensional neural space spanned by neurons recorded on *N* electrodes (*m<N*), we used the method described by Björck and Golub^73^. Consider the corresponding m-dimensional bases ***W***_*A*_ ∈ ℝ^*N*×*m*^ and ***W***_*B*_ ∈ ℝ^*N*×*m*^ provided by the m leading PCs/cPCs, we calculated their m by m inner product matrix 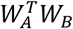, and performed a singular value decomposition to obtain

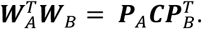

The matrices ***P***_*A*_ ∈ ℝ^*m*×*m*^ and ***P***_*B*_ ∈ ℝ^*m*×*m*^ were orthonormal vectors, which served as bases to transform the respective subspaces ***W***_*A*_ and ***W***_*B*_ onto a common subspace. The matrix *C* was a diagonal matrix whose elements were the ranked cosines of the principal angles *θ*_*i*_, *i* = 1, …, μ:

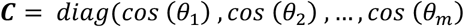

To assess whether the subspace angles between pairs of subspaces were significantly small, we compared them to the subspace angles between column-shuffled ***W***_*A*_ and ***W***_*B*_, which preserved the key statistics of the individual subspace^74^. We used the 0.1th percentile of the distributions of shuffled subspace angles to define a threshold below which angles can be considered significantly small (with a probability P < 0.001).

### Spike-triggered average analysis of CE HGA

To determine the relationship between spikes on all available electrodes and CE HGA with high temporal resolution, we used the spike times of all successful trials sampled at 1 kHz to perform spike-triggered average analysis. To compute instantaneous HGA, we used a 4^th^ order Butterworth band-pass filter at 200-300Hz bi-directionally on the raw LFP signals sampled at 1 kHz to remove any absolute time delay. Then we applied Hilbert transform on the filtered signals and squared the amplitude to compute the instantaneous HGA. This method enabled 1-ms resolution in the spike-triggered signal. For any electrode *n* and its spike event *i*_*n*_, we first segmented CE HGA from −50 ms to +70 ms relative to the spike time of *i*_*n*_ as *HGA*(*i*_*n*_) and averaged these CE HGA segments:

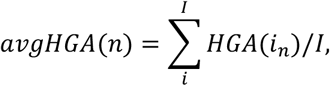

where *I* was the total number of spikes in the concatenated spiking activity recorded on the electrode *n*. These spike-triggered average segments were further standardized. We found the argmax of the *avgHGA*(*n*) to be the peak time *p* for any given electrode *n*. Finally, we reported the confidence interval from the distribution of *p(n)* to be the range of peak time of CE HGA triggered by the electrode.

### cPC projection of ONF and hand control data

To investigate how much the cPCs occupied the spaces spanned by hand control and ONF control data, we projected the hand control data and ONF data onto different cPCs. First, we concatenated the trials of population spiking activity during hand control and ONF control 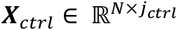 separately, where *ctrl* was either hand control or ONF control type and *j*_*ctrl*_ represented the length of concatenated trials of that control type, respectively. Then, we projected the Z-scored *X*_*ctrl*_ into the cPC space to obtain the m-dimensional latent 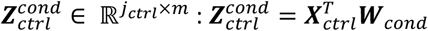, where *ccccnndd* represented either spike-HG angle of HG+ or SP+, and ***W***_*cond*_ ∈ ℝ^*N*×*m*^ represented the m dimensional space provided by the m leading cPCs. Finally, we calculated the variance accounted for (*VAF*) of each of the m dimensions of 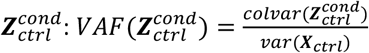, where *colvar* (∗) represented the variance calculation along the matrix column and *var*(∗) represented the variance calculated on the flattened matrix. We calculated the *VAF* for every *ctrl* - *cond* combination. For example, when *ctrl* was ONF control and *cond* was SP+, *VAF* was calculated on the trial-concatenated ONF control data projected into SP+ cPC space.

## Data availability

Data used for analysis to support the study results will be made publicly available upon publication at the Data Archive for the Brain Initiative repository.

## Code availability

Code related to contrastive principal analysis can be found at https://github.com/abidlabs/contrastive. Code used for analysis of the study data will be made publicly available upon publication on GitHub.

## Acknowledgements

We thank Zachary Wright for assistance with running experiments and technical support for the study, as well as Lee Miller for use of lab facilities and advice on the manuscript.

## References

1. Manning, J. R., Jacobs, J., Fried, I. & Kahana, M. J. Broadband Shifts in Local Field Potential Power Spectra Are Correlated with Single-Neuron Spiking in Humans. J. Neurosci. 29, 13613–13620 (2009).

2. Miller, K. J., Sorensen, L. B., Ojemann, J. G. & Den Nijs, M. Power-Law Scaling in the Brain Surface Electric Potential. PLoS Comput. Biol. 5, e1000609 (2009).

3. Hermes, D., Miller, K. J., Wandell, B. A. & Winawer, J. Stimulus Dependence of Gamma Oscillations in Human Visual Cortex. Cereb. Cortex 25, 2951–2959 (2015).

4. Nourski, K. V. et al. Temporal Envelope of Time-Compressed Speech Represented in the Human Auditory Cortex. J. Neurosci. 29, 15564–15574 (2009).

5. Crone, N. E., Miglioretti, D. L., Gordon, B. & Lesser, R. P. Functional mapping of human sensorimotor cortex with electrocorticographic spectral analysis. II. Event-related synchronization in the gamma band. Brain 121, 2301–2315 (1998).

6. Miller, K. J. et al. Spectral Changes in Cortical Surface Potentials during Motor Movement. J. Neurosci. 27, 2424–2432 (2007).

7. Flint, R. D. et al. The Representation of Finger Movement and Force in Human Motor and Premotor Cortices. eNeuro 7, ENEURO.0063-20.2020 (2020).

8. Ganguly, K., Khanna, P., Morecraft, R. J. & Lin, D. J. Modulation of neural co-firing to enhance network transmission and improve motor function after stroke. Neuron 110, 2363–2385 (2022).

9. Flint, R. D., Wright, Z. A., Scheid, M. R. & Slutzky, M. W. Long term, stable brain machine interface performance using local field potentials and multiunit spikes. J. Neural Eng. 10, 056005 (2013).

10. Silversmith, D. B. et al. Plug-and-play control of a brain–computer interface through neural map stabilization. Nat. Biotechnol. 39, 326–335 (2021).

11. Gilja, V. et al. Clinical translation of a high-performance neural prosthesis. Nat. Med. 21, 1142–1145 (2015).

12. Metzger, S. L. et al. Generalizable spelling using a speech neuroprosthesis in an individual with severe limb and vocal paralysis. Nat. Commun. 13, 6510 (2022).

13. Lachaux, J.-P., Axmacher, N., Mormann, F., Halgren, E. & Crone, N. E. High-frequency neural activity and human cognition: Past, present and possible future of intracranial EEG research. Prog. Neurobiol. 98, 279– 301 (2012).

14. Fries, P., Reynolds, J. H., Rorie, A. E. & Desimone, R. Modulation of Oscillatory Neuronal Synchronization by Selective Visual Attention. Science 291, 1560–1563 (2001).

15. Mendoza-Halliday, D. et al. A ubiquitous spectrolaminar motif of local field potential power across the primate cortex. Nat. Neurosci. 27, 547–560 (2024).

16. Rich, E. L. & Wallis, J. D. Spatiotemporal dynamics of information encoding revealed in orbitofrontal high-gamma. Nat. Commun. 8, 1139 (2017).

17. Canolty, R. T. et al. High Gamma Power Is Phase-Locked to Theta Oscillations in Human Neocortex. Science 313, 1626–1628 (2006).

18. Baratham, V. L. et al. Columnar Localization and Laminar Origin of Cortical Surface Electrical Potentials. J. Neurosci. 42, 3733–3748 (2022).

19. Pesaran, B., Pezaris, J. S., Sahani, M., Mitra, P. P. & Andersen, R. A. Temporal structure in neuronal activity during working memory in macaque parietal cortex. Nat. Neurosci. 5, 805–811 (2002).

20. Miller, K. J., Zanos, S., Fetz, E. E., Nijs, M.den & Ojemann, J. G. Decoupling the Cortical Power Spectrum Reveals Real-Time Representation of Individual Finger Movements in Humans. J. Neurosci. 29, 3132–3137 (2009).

21. Buzsáki, G. & Wang, X.-J. Mechanisms of Gamma Oscillations. Annu. Rev. Neurosci. 35, 203–225 (2012).

22. Crone, N. E., Sinai, A. & Korzeniewska, A. High-frequency gamma oscillations and human brain mapping with electrocorticography. in Progress in Brain Research vol. 159 275–295 (Elsevier, 2006).

23. Reimann, M. W. et al. A Biophysically Detailed Model of Neocortical Local Field Potentials Predicts the Critical Role of Active Membrane Currents. Neuron 79, 375–390 (2013).

24. Rule, M. E., Vargas-Irwin, C., Donoghue, J. P. & Truccolo, W. Contribution of LFP dynamics to single-neuron spiking variability in motor cortex during movement execution. Front. Syst. Neurosci. 9, (2015).

25. Gieselmann, M. A. & Thiele, A. Comparison of spatial integration and surround suppression characteristics in spiking activity and the local field potential in macaque V1. Eur. J. Neurosci. 28, 447–459 (2008).

26. Guyon, N. et al. Network Asynchrony Underlying Increased Broadband Gamma Power. J. Neurosci. 41, 2944–2963 (2021).

27. Sohal, V. S. & Rubenstein, J. L. R. Excitation-inhibition balance as a framework for investigating mechanisms in neuropsychiatric disorders. Mol. Psychiatry 24, 1248–1257 (2019).

28. Einevoll, G. T., Kayser, C., Logothetis, N. K. & Panzeri, S. Modelling and analysis of local field potentials for studying the function of cortical circuits. Nat. Rev. Neurosci. 14, 770–785 (2013).

29. Ray, S., Crone, N. E., Niebur, E., Franaszczuk, P. J. & Hsiao, S. S. Neural Correlates of High-Gamma Oscillations (60–200 Hz) in Macaque Local Field Potentials and Their Potential Implications in Electrocorticography. J. Neurosci. 28, 11526–11536 (2008).

30. Liu, J. & Newsome, W. T. Local field potential in cortical area MT: stimulus tuning and behavioral correlations. J. Neurosci. Off. J. Soc. Neurosci. 26, 7779–7790 (2006).

31. Ray, S. & Maunsell, J. H. R. Different Origins of Gamma Rhythm and High-Gamma Activity in Macaque Visual Cortex. PLoS Biol. 9, e1000610 (2011).

32. Belitski, A. et al. Low-Frequency Local Field Potentials and Spikes in Primary Visual Cortex Convey Independent Visual Information. J. Neurosci. 28, 5696–5709 (2008).

33. Buzsáki, G., Anastassiou, C. A. & Koch, C. The origin of extracellular fields and currents — EEG, ECoG, LFP and spikes. Nat. Rev. Neurosci. 13, 407–420 (2012).

34. Zanos, T. P., Mineault, P. J. & Pack, C. C. Removal of Spurious Correlations Between Spikes and Local Field Potentials. J. Neurophysiol. 105, 474–486 (2011).

35. Scheffer-Teixeira, R., Belchior, H., Leão, R. N., Ribeiro, S. & Tort, A. B. L. On High-Frequency Field Oscillations (>100 Hz) and the Spectral Leakage of Spiking Activity. J. Neurosci. 33, 1535–1539 (2013).

36. Bartoli, E. et al. Functionally Distinct Gamma Range Activity Revealed by Stimulus Tuning in Human Visual Cortex. Curr. Biol. 29, 3345-3358.e7 (2019).

37. Belluscio, M. A., Mizuseki, K., Schmidt, R., Kempter, R. & Buzsáki, G. Cross-Frequency Phase–Phase Coupling between Theta and Gamma Oscillations in the Hippocampus. J. Neurosci. 32, 423–435 (2012).

38. Gallego-Carracedo, C., Perich, M. G., Chowdhury, R. H., Miller, L. E. & Gallego, J.Á. Local field potentials reflect cortical population dynamics in a region-specific and frequency-dependent manner. eLife 11, e73155 (2022).

39. Leszczynski, M., Barczak, A. & Schroeder, C. Dissociation of broadband high-frequency activity and neuronal firing in the neocortex. Sci. Adv. 13 (2020).

40. Wright, Z. A., Rymer, W. Z. & Slutzky, M. W. Reducing Abnormal Muscle Coactivation After Stroke Using a Myoelectric-Computer Interface: A Pilot Study. Neurorehabil. Neural Repair 28, 443–451 (2014).

41. Hung, N.-T. et al. Wearable myoelectric interface enables high-dose, home-based training in severely impaired chronic stroke survivors. Ann. Clin. Transl. Neurol. 8, 1895–1905 (2021).

42. Mugler, E. M. et al. Myoelectric Computer Interface Training for Reducing Co-Activation and Enhancing Arm Movement in Chronic Stroke Survivors: A Randomized Trial. Neurorehabil. Neural Repair 33, 284–295 (2019).

43. Seo, G., Kishta, A., Mugler, E., Slutzky, M. W. & Roh, J. Myoelectric interface training enables targeted reduction in abnormal muscle co-activation. J. Neuroengineering Rehabil. 19, 67 (2022).

44. Khorasani, A. et al. Myoelectric interface for neurorehabilitation conditioning to reduce abnormal leg co-activation after stroke: a pilot study. J. NeuroEngineering Rehabil. 21, 11 (2024).

45. Matsumura, M., Chen, D., Sawaguchi, T., Kubota, K. & Fetz, E. E. Synaptic Interactions between Primate Precentral Cortex Neurons Revealed by Spike-Triggered Averaging of Intracellular Membrane Potentials In Vivo. J. Neurosci. 16, 7757–7767 (1996).

46. Barbieri, F., Mazzoni, A., Logothetis, N. K., Panzeri, S. & Brunel, N. Stimulus Dependence of Local Field Potential Spectra: Experiment versus Theory. J. Neurosci. 34, 14589–14605 (2014).

47. Bos, H., Diesmann, M. & Helias, M. Identifying Anatomical Origins of Coexisting Oscillations in the Cortical Microcircuit. PLOS Comput. Biol. 12, e1005132 (2016).

48. Francis, N. A., Elgueda, D., Englitz, B., Fritz, J. B. & Shamma, S. A. Laminar profile of task-related plasticity in ferret primary auditory cortex. Sci. Rep. 8, 16375 (2018).

49. Bartos, M., Vida, I. & Jonas, P. Synaptic mechanisms of synchronized gamma oscillations in inhibitory interneuron networks. Nat. Rev. Neurosci. 8, 45–56 (2007).

50. Kajikawa, Y. & Schroeder, C. E. How Local Is the Local Field Potential? Neuron 72, 847–858 (2011).

51. Lakatos, P., Karmos, G., Mehta, A. D., Ulbert, I. & Schroeder, C. E. Entrainment of Neuronal Oscillations as a Mechanism of Attentional Selection. Science 320, 110–113 (2008).

52. Sakata, S. & Harris, K. D. Laminar Structure of Spontaneous and Sensory-Evoked Population Activity in Auditory Cortex. Neuron 64, 404–418 (2009).

53. Łęski, S., Lindén, H., Tetzlaff, T., Pettersen, K. H. & Einevoll, G. T. Frequency Dependence of Signal Power and Spatial Reach of the Local Field Potential. PLoS Comput. Biol. 9, e1003137 (2013).

54. Ray, S., Hsiao, S. S., Crone, N. E., Franaszczuk, P. J. & Niebur, E. Effect of Stimulus Intensity on the Spike– Local Field Potential Relationship in the Secondary Somatosensory Cortex. J. Neurosci. 28, 7334–7343 (2008).

55. Ajiboye, A. B., Simeral, J. D., Donoghue, J. P., Hochberg, L. R. & Kirsch, R. F. Prediction of Imagined Single-Joint Movements in a Person With High-Level Tetraplegia. IEEE Trans. Biomed. Eng. 59, 2755–2765 (2012).

56. Milekovic, T. et al. Stable long-term BCI-enabled communication in ALS and locked-in syndrome using LFP signals. J. Neurophysiol. 120, 343–360 (2018).

57. Flint, R. D., Lindberg, E. W., Jordan, L. R., Miller, L. E. & Slutzky, M. W. Accurate decoding of reaching movements from field potentials in the absence of spikes. J. Neural Eng. 9, 046006 (2012).

58. Okun, M. et al. Diverse coupling of neurons to populations in sensory cortex. Nature 521, 511–515 (2015).

59. Ray, S., Niebur, E., Hsiao, S. S., Sinai, A. & Crone, N. E. High-frequency gamma activity (80–150Hz) is increased in human cortex during selective attention. Clin. Neurophysiol. 119, 116–133 (2008).

60. Kreiman, G. et al. Object Selectivity of Local Field Potentials and Spikes in the Macaque Inferior Temporal Cortex. Neuron 49, 433–445 (2006).

61. Henze, D. A. et al. Intracellular Features Predicted by Extracellular Recordings in the Hippocampus In Vivo. J. Neurophysiol. 84, 390–400 (2000).

62. Barrese, J. C. et al. Failure mode analysis of silicon-based intracortical microelectrode arrays in non-human primates. J. Neural Eng. 10, 066014 (2013).

63. Kozai, T. D. Y., Jaquins-Gerstl, A. S., Vazquez, A. L., Michael, A. C. & Cui, X. T. Brain tissue responses to neural implants impact signal sensitivity and intervention strategies. ACS Chem. Neurosci. 6, 48–67 (2015).

64. Prasad, A. et al. Comprehensive characterization and failure modes of tungsten microwire arrays in chronic neural implants. J. Neural Eng. 9, 056015 (2012).

65. McConnell, G. C. et al. Implanted neural electrodes cause chronic, local inflammation that is correlated with local neurodegeneration. J. Neural Eng. 6, 056003 (2009).

66. Flint, R. D., Scheid, M. R., Wright, Z. A., Solla, S. A. & Slutzky, M. W. Long-Term Stability of Motor Cortical Activity: Implications for Brain Machine Interfaces and Optimal Feedback Control. J. Neurosci. 36, 3623– 3632 (2016).

67. Karpowicz, B. M. et al. Stabilizing brain-computer interfaces through alignment of latent dynamics. 2022.04.06.487388 Preprint at 10.1101/2022.04.06.487388 (2022).

68. Fries, P. Neuronal Gamma-Band Synchronization as a Fundamental Process in Cortical Computation. Annu. Rev. Neurosci. 32, 209–224 (2009).

69. Lee, K.-H., Williams, L. M., Breakspear, M. & Gordon, E. Synchronous Gamma activity: a review and contribution to an integrative neuroscience model of schizophrenia. Brain Res. Rev. 41, 57–78 (2003).

70. Herrmann, C. S., Fründ, I. & Lenz, D. Human gamma-band activity: A review on cognitive and behavioral correlates and network models. Neurosci. Biobehav. Rev. 34, 981–992 (2010).

71. Bédard, C., Kröger, H. & Destexhe, A. Modeling Extracellular Field Potentials and the Frequency-Filtering Properties of Extracellular Space. Biophys. J. 86, 1829–1842 (2004).

72. Abid, A., Zhang, M. J., Bagaria, V. K. & Zou, J. Exploring patterns enriched in a dataset with contrastive principal component analysis. Nat. Commun. 9, 2134 (2018).

73. Bjorck, A. & Golub, G. H. Numerical Methods for Computing Angles Between Linear Subspaces. Math. Comput. 27, 579 (1973).

74. Gallego, J. A. et al. Cortical population activity within a preserved neural manifold underlies multiple motor behaviors. Nat. Commun. 9, 4233 (2018).

