## Supplementarty figures for "Active Dissociation of Intracortical Spiking and High Gamma Activity"

### Extended Data

**Extended Data Table 1**. Number of sessions of recording for each control electrode (CE)

| Monkey-CE | C. CE 10 | C. CE 63 | C. CE 81 | J. CE 29 | J. CE 30 | J. CE 52 | M.CE 66 | M. CE 73 | M. CE 92 |
|---|---|---|---|---|---|---|---|---|---|
| Number of sessions | 15 | 5 | 5 | 6 | 4 | 4 | 7 | 10 | 5 |

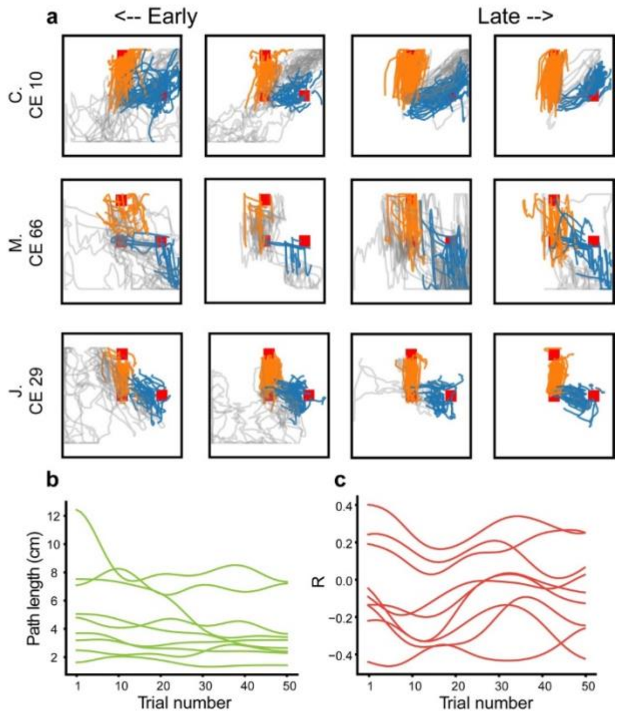


**Extended Data Figure 1**. Monkeys quickly became proficient at the ONF task. **a**, Example cursor traces of the last 2 s of all trials in 4 sessions for three example monkey-CE combinations. Row: each monkey-CE combination. Column: sessions. **b**, Path length of the first 50 trials (the minimum number of trials across the monkey-CE combinations within the first two sessions) for each monkey-CE combination showed significant decrease. ($p<0.05$, one sample t test) **c**, R between spike rate and HGA of the first 50 trials for each monkey-CE combination. Six out of 9 CEs had R of 0 or less from the beginning of the first two sessions, and 8 of 9 had R less than 0.25 to start with. Data in **b** and **c** were smoothed by a Gaussian kernel with $\sigma$=5.

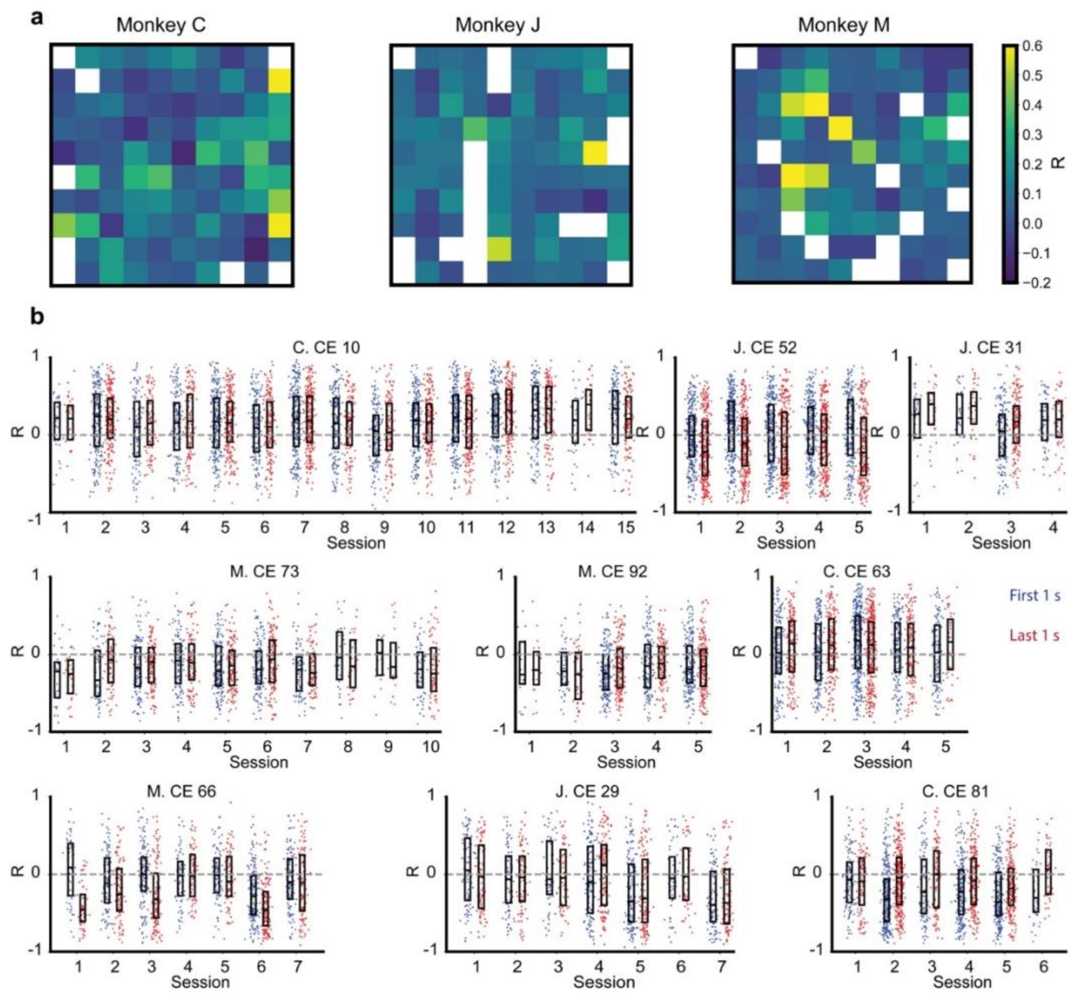


**Extended Data Figure 2.** Correlation between HGA, spike rate, was moderate across electrodes and did not demonstrate significant change across sessions on the CE. **a**, Mean correlation (R; see Methods) between single-trial spike rate and HGA recorded on the same electrodes of the last second preceding reward during hand control for each monkey (monkey C: 32 sessions, monkey J: 6 sessions, monkey M: 6 sessions). **b**, Correlation coefficient between spike rate and HGA on the CE of the first (blue) and last (red) 1 s of all trials (each dot is one trial), including failed trials, over sessions.

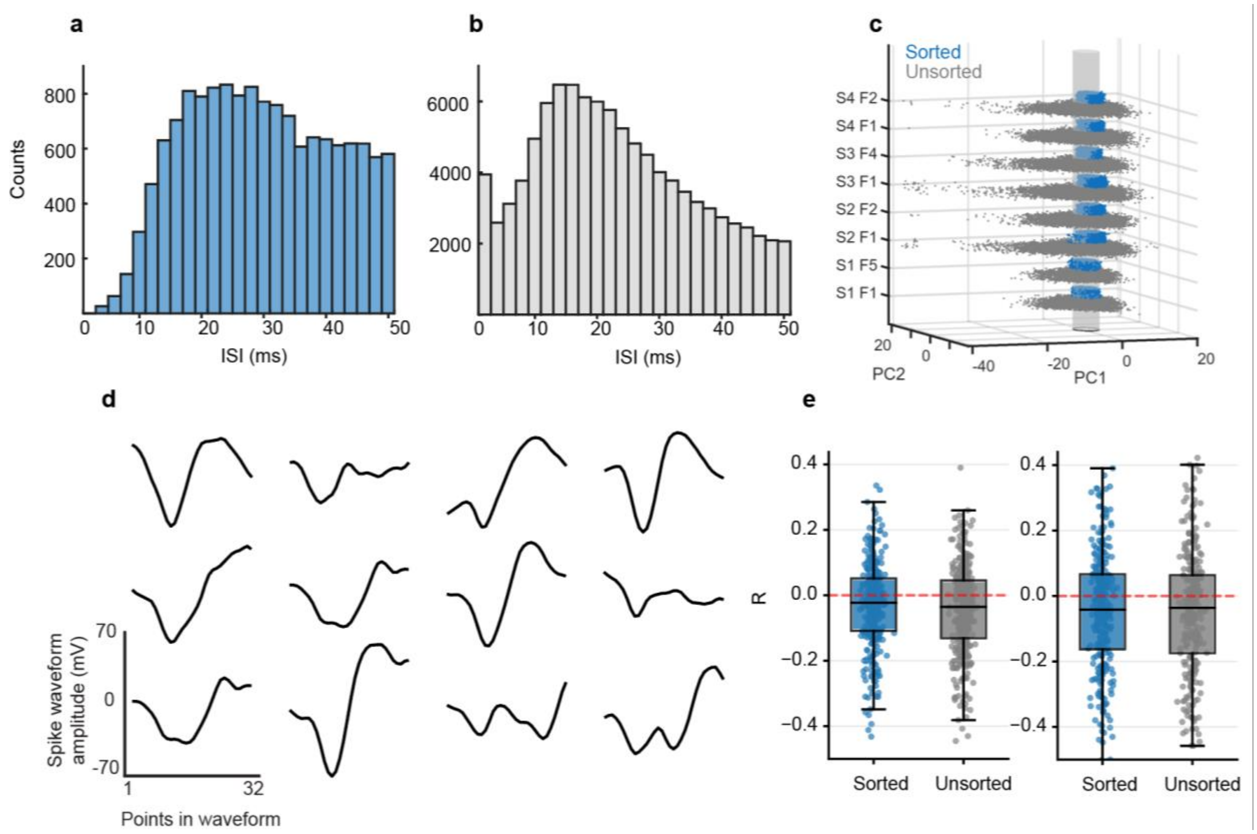


**Extended Data Figure 3.** Examples of inter-spike intervals (ISIs), waveforms and their clusters, as well as correlation (R) between CE HGA and sorted spikes when using a 3-s.d. threshold. **a**, **b**, ISIs from sorted (**a**; blue) and unsorted (**b**; gray) spike waveforms in J-CE 52 (8 files over 4 sessions). The unsorted waveforms have a relatively high percentage of very short ISIs, indicating multiple (likely many) units. **c,** Principal component projections (PC1 and PC2) of sorted (blue) and unsorted (grey) spikes on the J-CE 52. Axes are in units of s.d. of each PC. The vertical axis indicated the session-file indices. The first and last files (F) of each session (S) are shown. For visualization, the vertical values of the points were randomly jittered in each cloud, which would otherwise occupy a 2D region for each file. Translucent cylinder represents 2 s.d. in each direction of the sorted spikes in session 1, file 1 (S1 F1). **d**, Twelve examples of waveforms sampled from the PC space in one file. **e**, Correlation (R) between CE HGA and sorted spikes (blue) or unsorted spikes (grey) over 4 files in 2 sessions in J CE 29 (left) and 8 files in 4 sessions in J CE 52 (right). Each dot represents an R value of the last 1 s of a trial. Both sorted and unsorted spikes were uncorrelated with CE HGA.

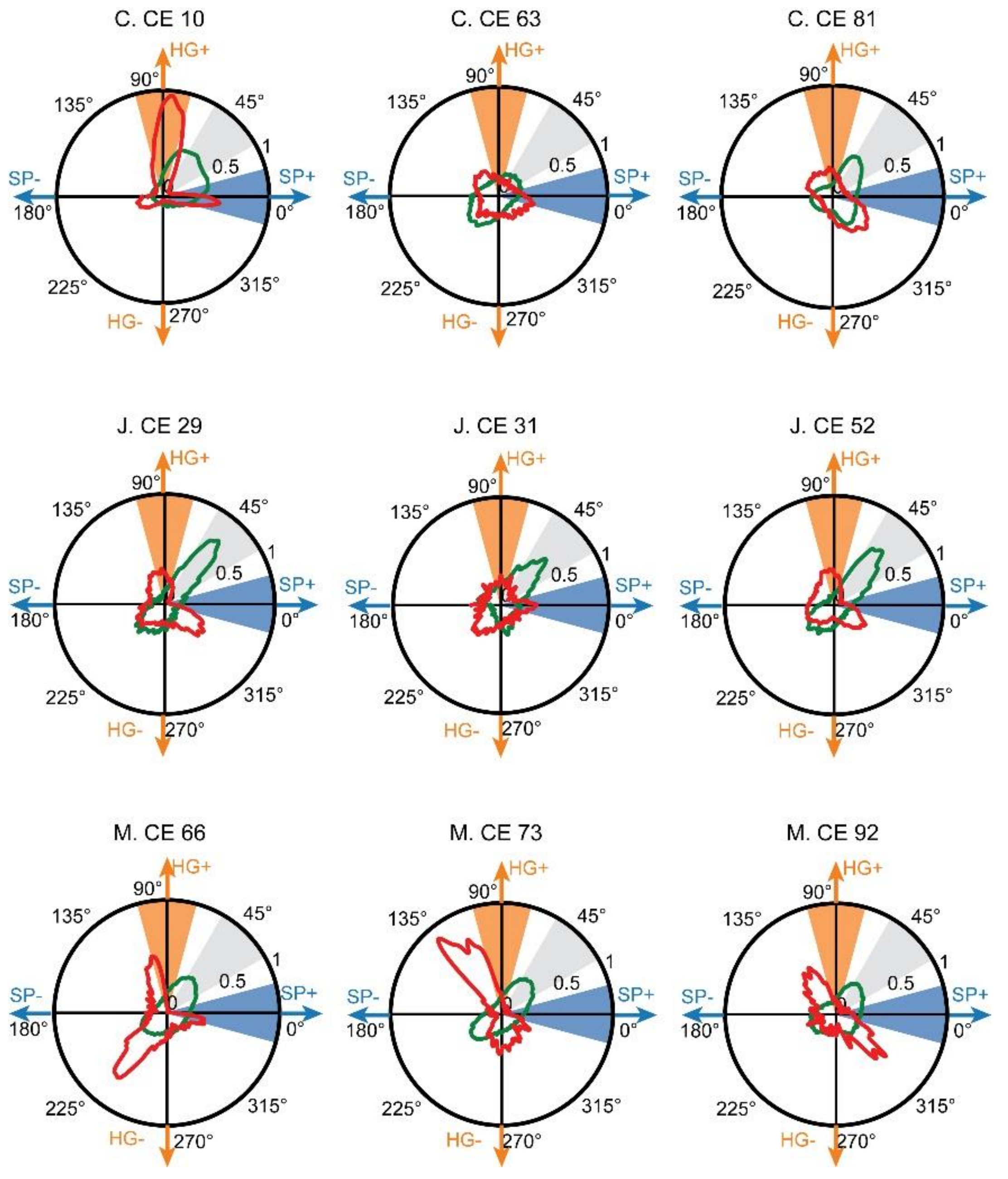


**Extended Data Figure 4**. Polar plot of the 2D distribution of mean spike-HG angles of each 50-ms bin in the 1 second preceding the reward time of ONF files (red) and hand control files (green) from the learned sessions of all monkey-CE combinations. Orange sector: HG+ state; blue sector: SP+ state. Gray sector: state at which both signals had high modulation. Note that states in the 135° and 315° directions also corresponded to independent modulation of HGA and spiking. States near 225° corresponded to times at which there was minimal spiking and HGA, likely times when the animal was not actively engaged in the task

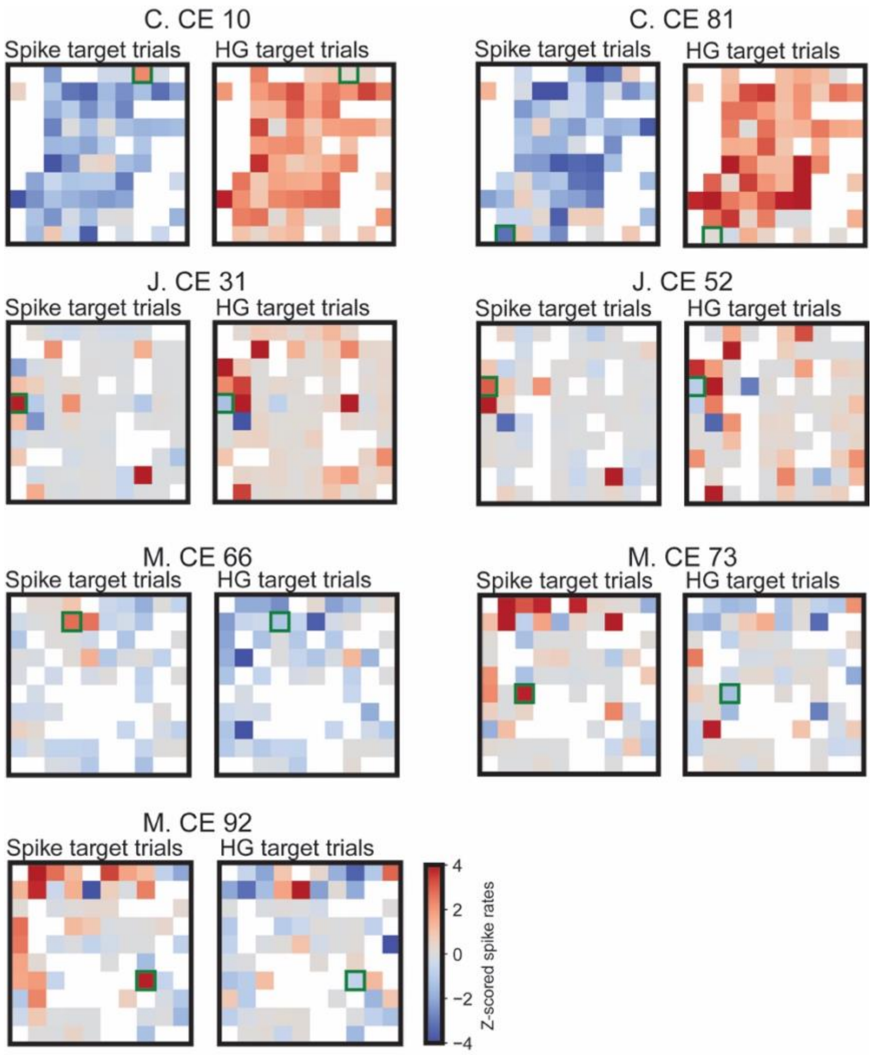


**Extended Data Figure 5.** Trial-averaged, z-scored spike rates across the array over the last second of spike target trials (left plots) and HG target trials (right plots) during ONF control in all monkey-CE combinations. White squares denote shunted electrodes. CEs are outlined in green.

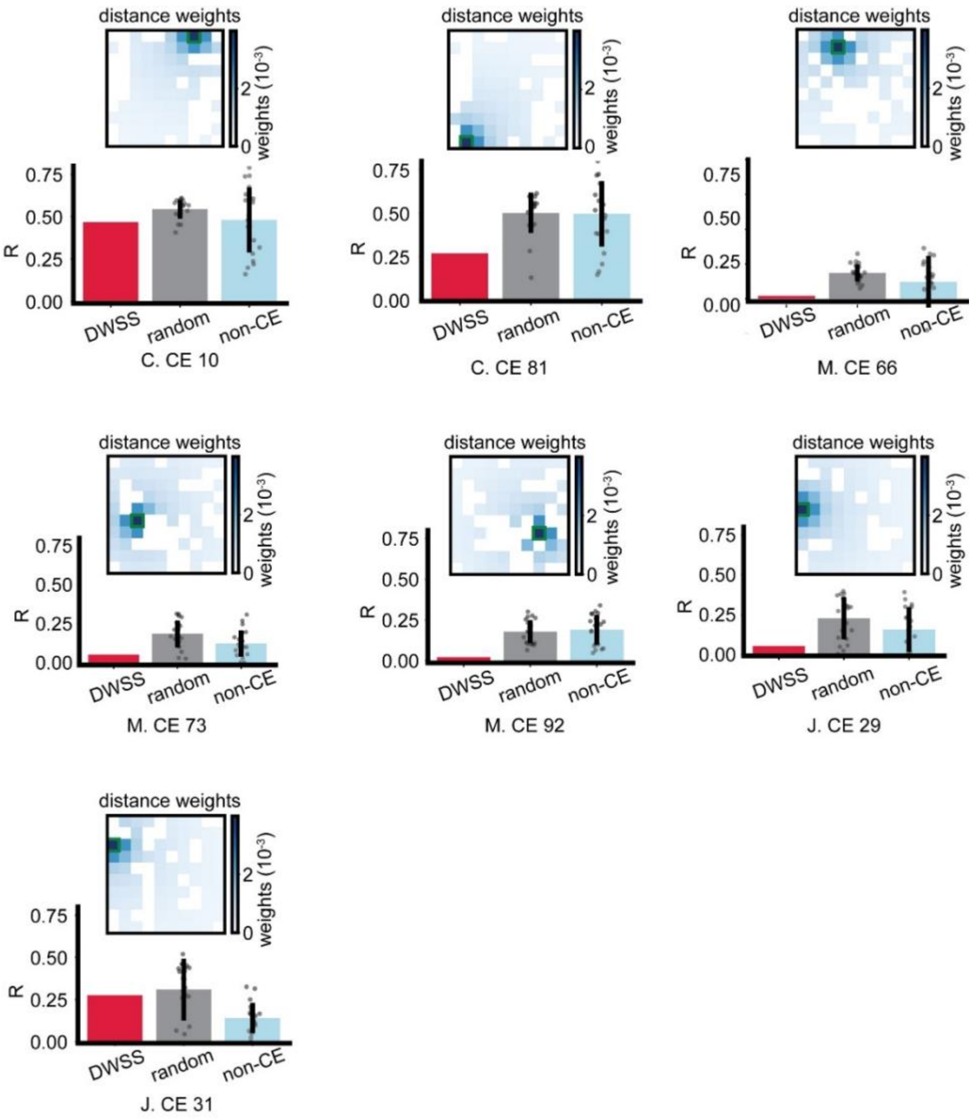


**Extended Data Figure 6.** Correlation coefficient (R) between CE HGA and the control-electrode distance-weighted sum of spikes (CE DWSS, red bar), the mean (±s.d.) R between CE HGA and DWSS of 20 randomly selected electrodes (rand DWSS, grey bar), the mean (±s.d.) R between the HGA on 20 randomly selected non-CE electrodes and DWSS of each of those non-CE electrodes (random, gray bar; each dot is one R with one electrode), as well as the mean (±s.d.) R between HGA and DWSS pairs from 20 randomly selected electrodes excluding the CE (non-CE, light blue bar). The R values between CE HGA and CE DWSS were not significantly larger than those between CE HGA and random DWSS, or than those between the non-CE HGA and DWSS (n.s., one- sample, one-sided Mann-Whitney U test). Inset, the weights used to compute the CE DWSS. Plots include all monkey-CE combinations except those in Fig. 5g.

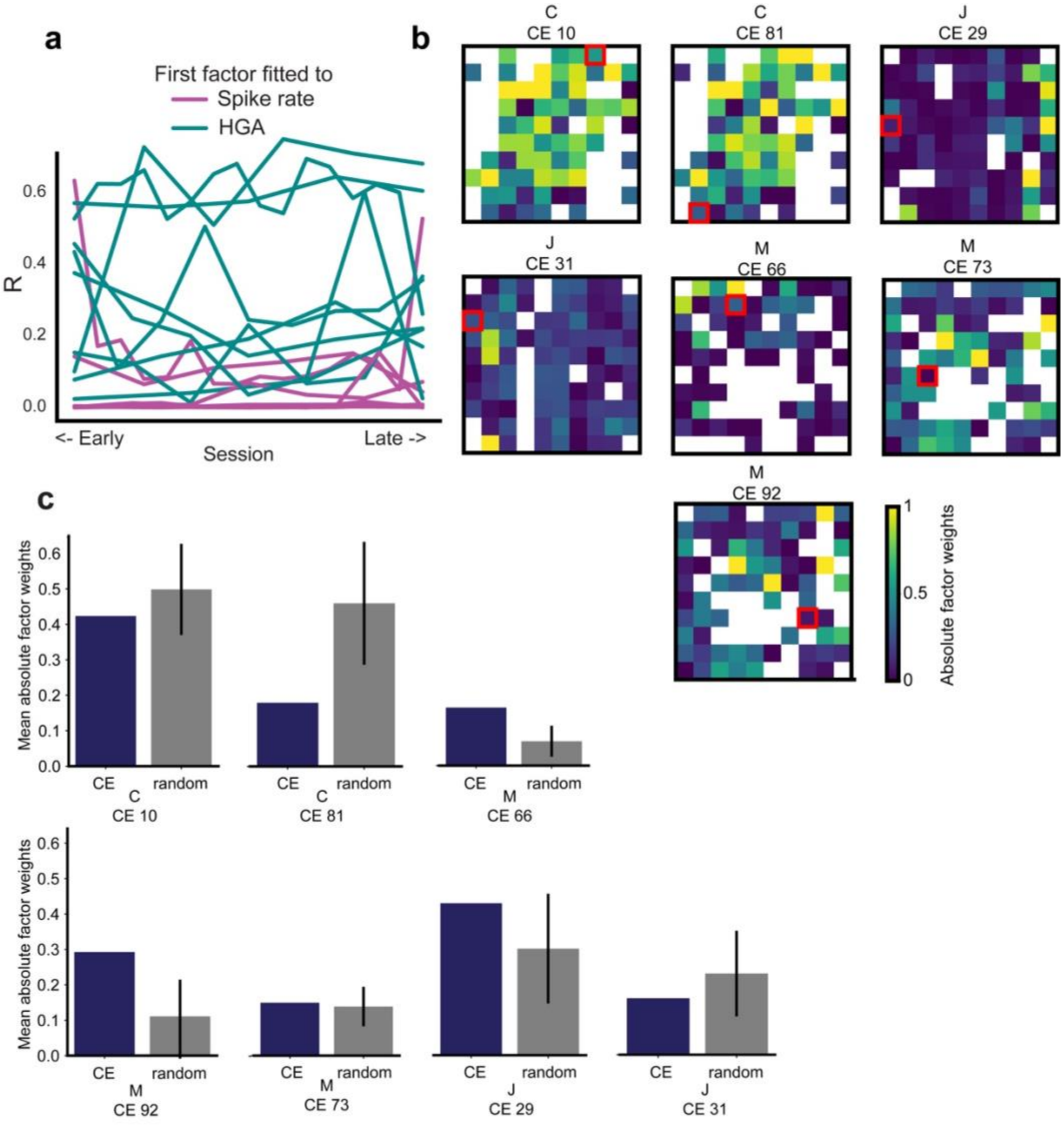


**Extended Data Figure 7. a**, Correlation (R) of MLR fit from the first co-firing pattern to spike rate (magenta), or to the HGA (turquoise) by sessions over all CEs. The factor weights were calculated using all the sessions combined and then were used to project in each session separately to generate the first factor. We did not observe any increase or decrease of the correlation across sessions ($p>0.9$, one sample t-test). **b**, Absolute values of the factor (co-firing) weights of all monkey-CE combinations except the two examples shown in Fig. 6d. CEs are outlined in red. White electrodes represent either shunted or those that did not exceed the minimum firing rate (see Methods). **c**, The mean value of the absolute co-firing weights on the electrodes that are adjacent to the CE (dark blue), as well as the mean (±s.d.) of average value of the absolute co-firing weights on the electrodes that are adjacent to 20 randomly selected electrodes (gray) of all monkey-CE combinations except the two examples shown in Fig. 6e. These R values were not significantly different for any CE (n.s., one sample one-sided Mann-Whitney U test).

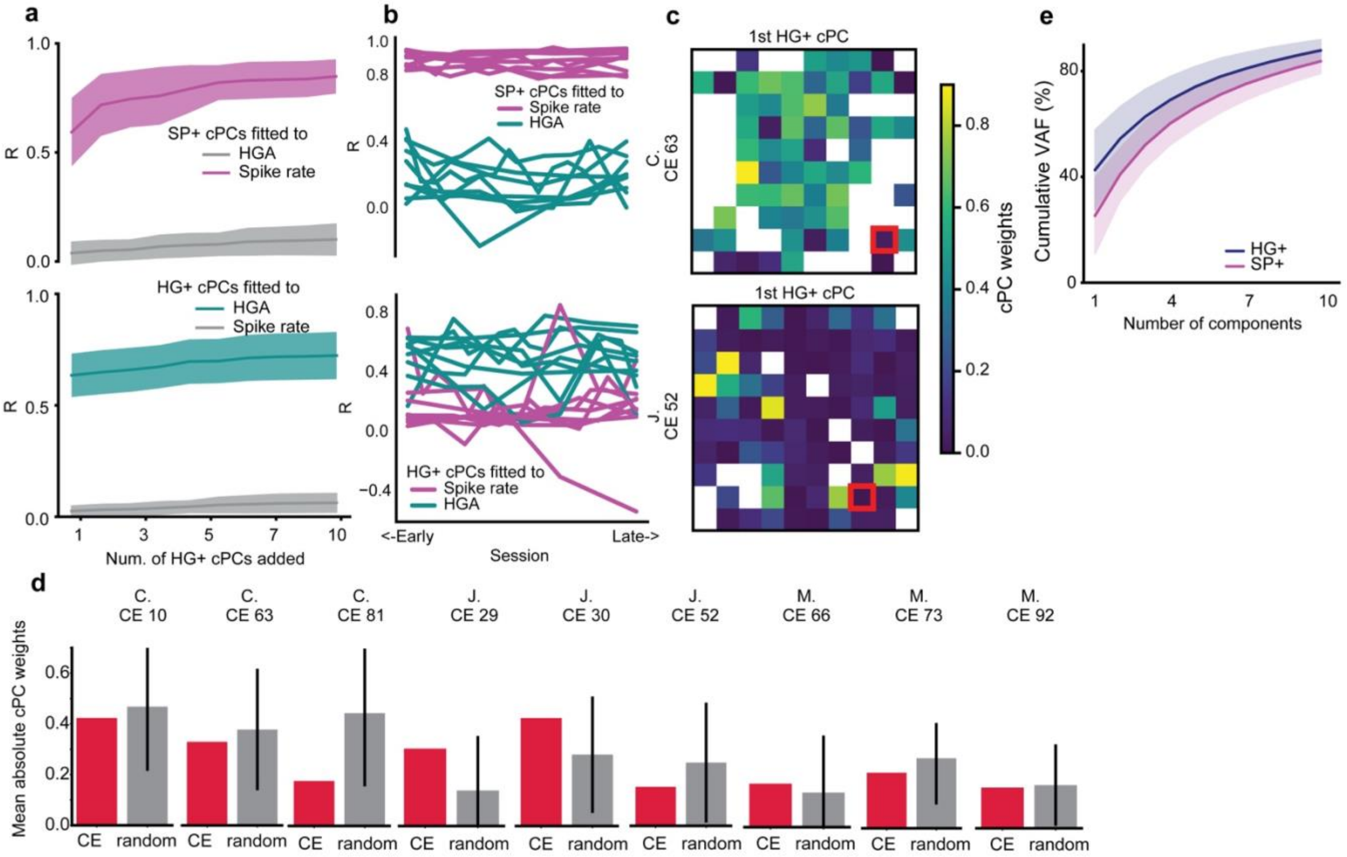


**Extended Data Figure 8.** Spike rate and HGA modulation on the CE involve activity in different subspaces of the neuronal ensemble. **a**, Top, correlation (R) of the linear fit of ensemble SP+ cPC projections to CE spike rate (magenta) and CE HGA (gray) separately. Bottom, fit of ensemble HG+ cPC projections to CE spike rate (gray) and CE HGA (turquoise) separately on the CE with increase of number of cPCs from 1-10. **b**, Top: correlation (R) of MLR fit of top 10 SP+ cPC projections to CE spike rate (magenta) and CE HGA (turquoise) by sessions over all CEs. Bottom: fit of top 10 HG+ cPC projections to CE spike rate (magenta) and CE HGA (turquoise) by sessions over all CEs. The cPC weights were calculated using all the sessions combined and then were projected to each session separately to generate the cPC projections. We did not observe any increase or decrease of R across sessions ($p>0.9$, one sample t-test).**c** , Weights of the first HG+ cPC across the array for one CE (red outline) in Monkey C (top) and Monkey J (bottom). **d**, The mean value of the absolute first HG+ cPC weights on the 4 electrodes that are adjacent to the CE (red), as well as the mean (±s.d.) of mean value of 4 randomly selected electrodes with 1000 repetitions (gray) of all monkey-CE combinations. The CE mean was not significantly larger than the mean of the randomly selected electrodes in any case ($p>0.05$, one-sided permutation test). **e**, Cumulative variance accounted for (VAF) for components constructed by singular value decomposition of SP+ data (magenta) and HG+ data (blue). Shaded area: s.d..

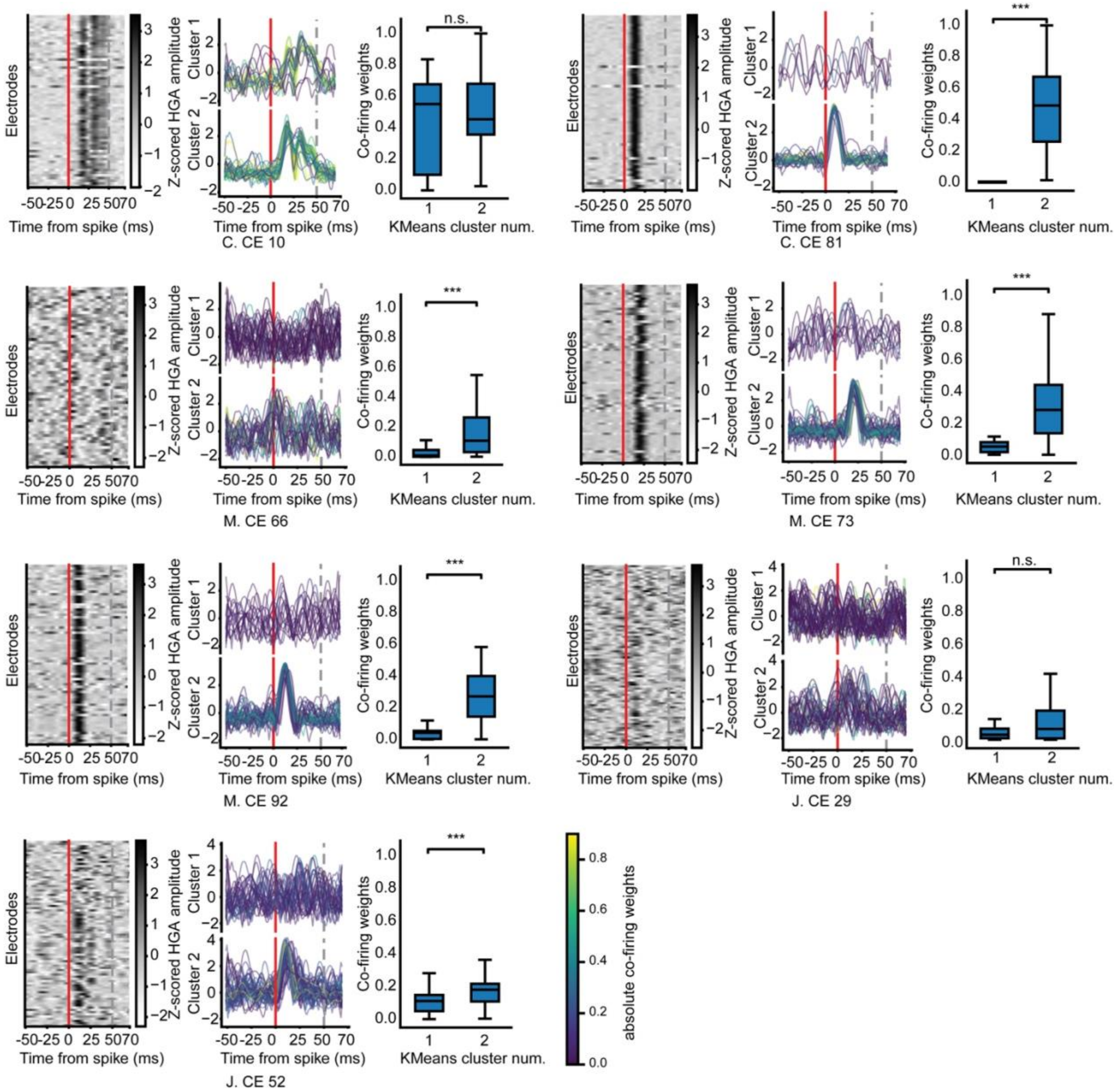


**Extended Data Figure 9.** Spikes contributing most to co-firing also contributed most to HGA for the remainder of the monkey-CE combinations not shown in Fig. 7. First and fourth column, spike-triggered averaged HGA of the CE showed a consistent activity pattern when aligned to the spike time of other electrodes. Second and fifth column, K-means clustering results using 2 clusters on the STA HGA traces, color-coded by the absolute co-firing weights. These clusters correspond generally to the "no response" and "with response" clusters in Fig. 7d. Third and sixth column, a comparison of the absolute co-firing weights of the two clusters. (***p<0.005, Wilcoxon signed-rank test).

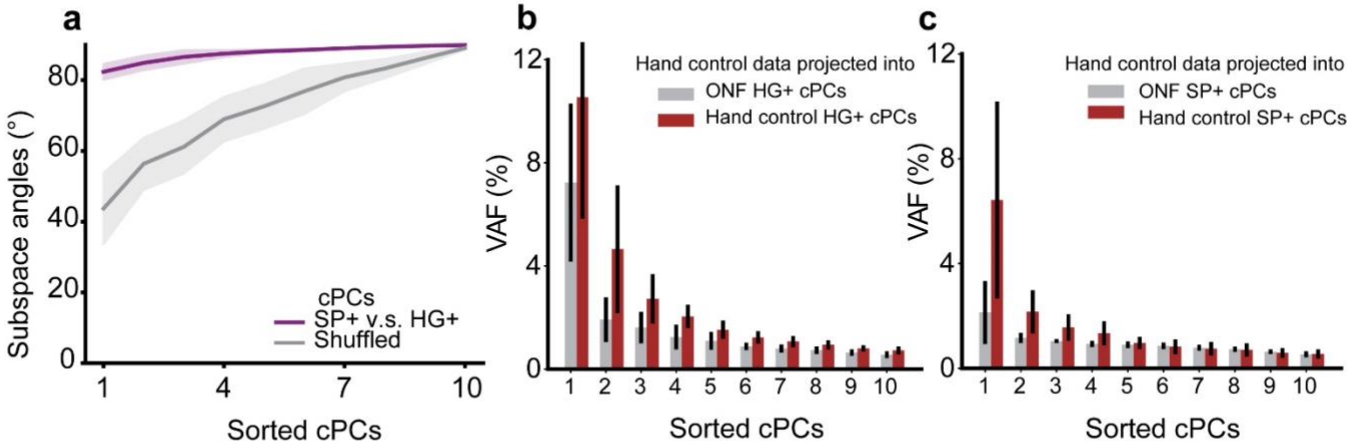


**Extended Data Figure 10.** Subspace analysis of cPCs. **a**, Subspace angles between the top ten cPCs (sorted by variance accounted for) of SP+ data and HG+ data. Leading subspace angles were significantly different from chance ($p<0.001$; see Methods). Grey lines and shaded areas indicate subspace angles calculated using shuffled subspaces. **b**, **c**, VAF when hand control data were projected into the subspaces defined by the top ten **b,** HG+ cPCs computed with ONF data (grey) or hand control data (brown), and **c,** SP+ cPCs computed with ONF data (grey) or hand control (brown). Error bars indicated s.d.
